# SOMATODENDRITIC VASOPRESSIN RELEASE COUPLES INTRINSIC EXCITABILITY TO POPULATION-LEVEL ACTIVITY

**DOI:** 10.64898/2026.09.22.753511

**Authors:** Matthew K. Kirchner, Fiona C. Crane, Ismail A. Ahmed, Javier E. Stern

## Abstract

The transition from single-neuron excitability to organized population firing is a fundamental feature of neural circuits, yet the mechanisms that govern this transition remain poorly understood. Magnocellular vasopressin (VP) and oxytocin neurons are intermingled within the hypothalamus, yet display strikingly different population dynamics, with VP output sustained by a distributed, largely asynchronous activity. Here, we identify a local mechanism by which somatodendritically released VP acts as a diffusible signal that converts an intrinsic excitability mechanism into spatially and temporally organized feedback across the VP population. Activity-dependent somatodendritic VP release recruited a rapid autocrine feedback that transiently potentiated the slow afterhyperpolarization (sAHP) through V1aR signaling which in turn strengthened spike-frequency adaptation and restrained firing. The same signal also acted in a diffusible, paracrine manner on neighboring VP neurons, producing distance- and time-dependent sAHP modulation, with rapid potentiation at short range and delayed inhibition at greater distances. Focal VP uncaging further revealed preferential engagement of somatic over dendritic signaling. Together, these findings show how a diffusible neuropeptide can convert an intrinsic excitability mechanism into spatially structured intercellular feedback, providing a candidate mechanism for sustaining distributed, asynchronous VP population output during prolonged homeostatic demand.

## INTRODUCTION

Neuronal activity in the central nervous system is shaped by intrinsic membrane properties, synaptic inputs, and slower modulatory signals such as neuropeptides^1,2^. Yet neurons rarely operate in isolation: adaptive circuit output also depends on mechanisms that distribute and coordinate activity across populations of cells. How such population-level organization emerges from local intrinsic and neuromodulatory processes remains poorly understood.

Hypothalamic vasopressin (VP) neurons provide a powerful model in which to address this question. These neurons combine well-defined intrinsic firing mechanisms with activity-dependent somatodendritic peptide release capable of acting locally on both the releasing neuron and neighboring cells^3–5^. As a result, VP neurons offer a unique opportunity to examine how mechanisms governing the excitability of individual neurons intersect with diffusible neuropeptide signaling that may organize activity across a neuronal population.

Magnocellular VP neurons located in the supraoptic (SON) and paraventricular (PVN) nuclei are essential for osmotic, cardiovascular, and fluid homeostasis^6,7^. Their systemic output depends on activity-dependent release of VP from axon terminals in the posterior pituitary, where action potential pattern strongly influences hormone secretion^8–11^. In individual VP neurons, phasic firing is particularly effective at promoting axonal peptide release, and substantial work has defined the synaptic and intrinsic mechanisms that contribute to this patterning^8,12–15^. At the population level, however, the homeostatic VP response to an osmotic challenge depends on more than the firing properties of any single neuron. Unlike oxytocin neurons, whose coordinated burst firing supports pulsatile secretion^16–19^, VP neurons must sustain a more distributed and largely asynchronous population output during osmotic challenge, such that circulating VP reflects the integrated activity of many neurons firing asynchronously over time^9,14,20^. Still, the precise mechanism by which this population-level organization is stabilized and adjusted remains unresolved.

The firing pattern of an individual VP neuron reflects the interaction between synaptic inputs and intrinsic membrane conductances. Among the intrinsic mechanisms that shape excitability, afterpotentials are especially important because they regulate the probability, frequency, and temporal structure of subsequent spiking^21,22^. In magnocellular neurons, both depolarizing and hyperpolarizing afterpotentials contribute to activity patterning, but slow afterhyperpolarizations (sAHPs) are particularly well positioned to constrain repetitive firing by promoting spike frequency adaptation over extended timescales^4^. These intrinsic inhibitory mechanisms are therefore well positioned to couple a neuron’s recent activity to its subsequent output.

In addition to their endocrine secretion from axon terminals, VP neurons also release their peptidergic cargo locally from somatic and dendritic compartments^23–25^. Somatodendritic release is activity-dependent and has long been implicated in feedback regulation within the magnocellular system. Previous studies suggest that locally released VP can restrain neuronal firing through V1a receptor (V1aR) signaling^26,27^, supporting the idea that VP-SDR acts as an autocrine inhibitory signal. Local co-released dynorphin was shown to inhibit the depolarizing afterpotential, producing burst termination^28–30^. Moreover, we previously showed that somatodendritically released VP diffuses through the extracellular space far enough to influence neighboring neurons locally and to function over longer ranges as an intra- and interpopulation signal^31,32^. This raises the possibility that VP-SDR could serve not only as a single-cell feedback signal, but also as a local mechanism for distributing excitability across the VP neuronal population.

Despite these observations, the cellular mechanisms through which VP-SDR feeds back onto VP neurons remain unclear. Specifically, it is not known whether VP acts by engaging intrinsic mechanisms such as the sAHP, whether this regulation can be recruited by endogenous activity-dependent peptide release, or whether the same signal extends beyond the releasing neuron to influence neighboring VP neurons in a spatially organized manner. Here, we show that somatodendritically released VP diffuses locally to engage autocrine and paracrine slow afterhyperpolarizing feedback, providing a mechanism that couples intrinsic excitability to the spatiotemporal organization of activity across the magnocellular VP network.

## RESULTS

### VP enhances the integrated sAHP and spike frequency adaptation in vasopressin neurons

VP neurons were targeted using a transgenic VP-GFP reporter rat^33^. During current-clamp recordings, sAHPs were isolated using a pharmacological cocktail. Because other afterpotentials such as the slow depolarizing afterpotential (DAP) and mAHP overlap with the time course of the sAHP, we included Cs^+^ to block the sDAP^34^, and apamin to block the mAHP^12,35^ to fully unmask the sAHP (see methods). Additionally, previous studies have shown that VP can mediate an inhibitory effect synaptically by reducing excitatory transmission^36^. We therefore included DNQX, AP5, and picrotoxin to eliminate synaptic influences on sAHP properties.

Bath-applied VP ([Arg^8^]-Vasopressin) enhanced the area but not the amplitude of the sAHP evoked by a fixed train of action potentials (**Fig. 1A**). Similar effects were observed when sAHPs were evoked by step depolarizations **(Supplementary Fig. 1A)**. Measurements of intracellular Ca^2+^ levels during the train of action potentials revealed that VP induced enhancements in Ca^2+^ signal amplitude, area, and decay tau during the pulse (**Fig. 1B**). Consistent with previous studies, VP application increased the baseline Ca^2+^signal fluorescence^37,38^. VP effects on the sAHP translated into changes in spiking activity, as the number of action potentials evoked during a depolarizing step was significantly reduced following bath-applied VP (**Fig. 1C**). This also reflected as an enhanced spike frequency adaptation (SFA) throughout the spike train. We constructed plots of instantaneous firing frequency as a function of spike time during the depolarizing steps, and the data were fitted with a single exponential function to estimate the decay time constant (adaptation tau)^39^. Our results show that the SFA adaptation tau was significantly reduced after VP bath application (**Fig. 1D**). Together, these results indicate that VP increases the integrated sAHPs and the associated SFA, leading to an overall prolonged intrinsic inhibitory state in VP neurons.

**Figure 1.**
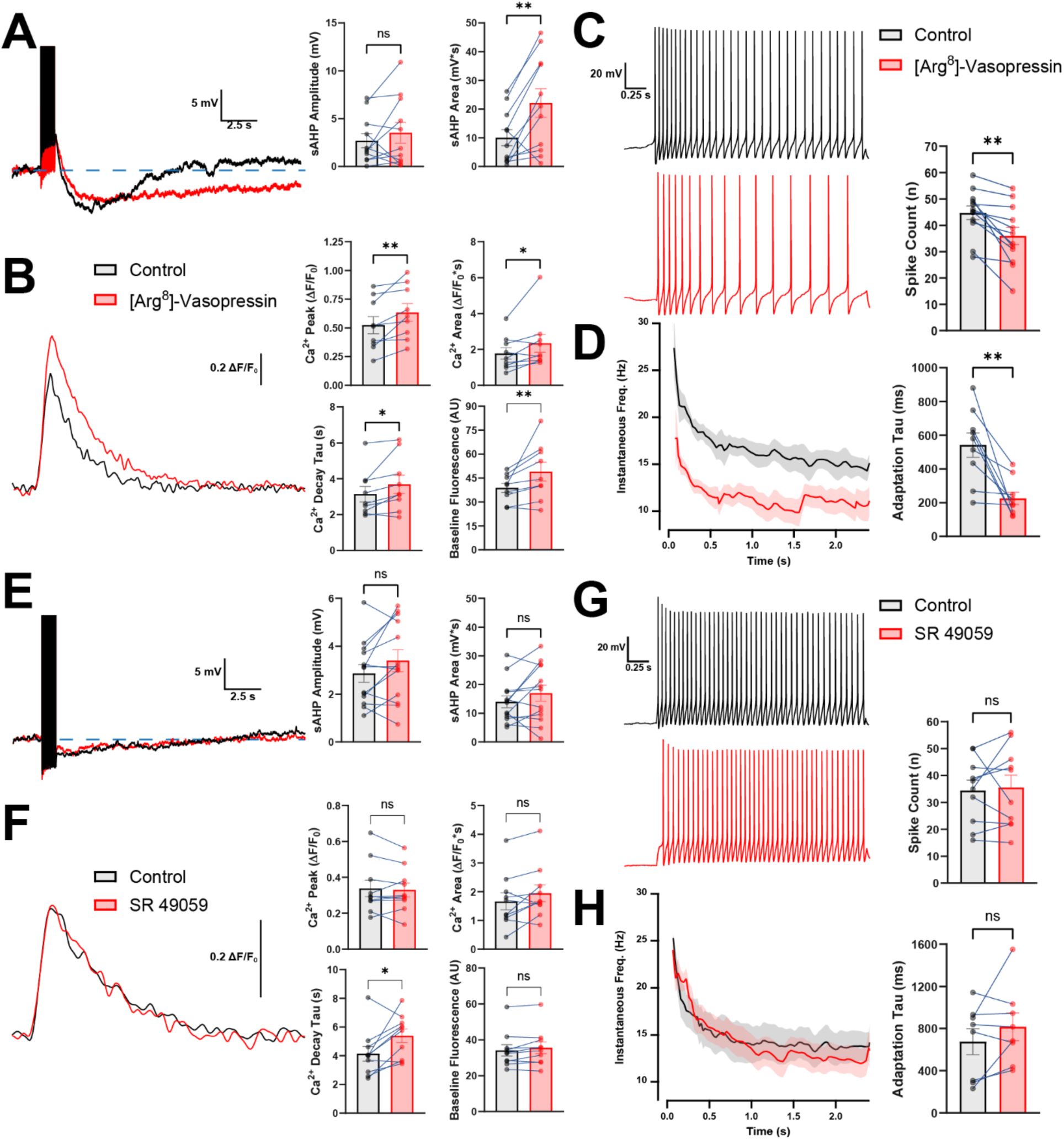
VP enhances sAHP magnitude but is not required for its activation. **(A)** Exogenous bath application of [Arg^8^]-Vasopressin enhances area (n = 11, paired t-test, t(10) = 3.584, P = 0.0050) but not amplitude (*n =* 11, paired t-test, *t*(10) = 1.072, *P =* 0.3089) of sAHPs evoked by a fixed train of spikes. **(B)** [Arg^8^]-Vasopressin enhances corresponding Ca^2+^ signals, demonstrating increased Ca^2+^ peak (*n =* 9, paired t-test, *t*(8) = 3.392, *P =* 0.0095), area (*n =* 9, Wilcoxon test, *P =* 0.0095), decay tau (*n =* 9, paired t-test, *t*(8) = 2.520, *P =* 0.0358), and baseline fluorescence (*n =* 9, paired t-test, *t*(8) = 2.476, *P =* 0.0383). **(C)** Spike trains evoked by a single 50 pA depolarizing current step demonstrate lower spike count after exogenous [Arg^8^]-Vasopressin (*n =* 12, paired t-test, *t*(11) = 4.110, *P =* 0.0020). **(D)** Instantaneous frequency plotted as a function of time reveals dramatic reduction of spike activity and enhancement of spike frequency adaptation. Consequently, adaptation taus are also significantly smaller after [Arg^8^]-Vasopressin compared to controls (*n =* 9, paired t-test, *t*(8) = 3.908, *P =* 0.0045). **(E)** Exogenous bath application of V1aR antagonist SR 49059 failed to affect amplitude (*n =* 13, paired t-test, *t*(12) = 1.911, *P =* 0.0801) or area (*n =* 13, paired t-test, *t*(12) = 1.541, *P =* 0.1493) of sAHPs evoked by a fixed train of spikes. **(F)** SR 49059 fails to affect corresponding Ca^2+^ peak (*n =* 10, paired t-test, *t*(9) = 0.5471, *P =* 0.5976), area (*n =* 10, paired t-test, *t*(9) = 1.981, *P =* 0.0790), and baseline (*n =* 10, Wilcoxon test, *P =* 0.0644), but did affect the decay tau (*n =* 10, paired t-test, *t*(9) = 2.839, *P =* 0.0194). **(G)** Spike trains evoked by a single depolarizing current step are unaffected by exogenous SR 49059 (*n =* 10, paired t-test, *t*(9) = 0.3322, *P =* 0.7474). **(H)** Instantaneous frequency plotted as a function of time reveals no change in spike activity nor enhancement of spike frequency adaptation. Adaptation taus are also unaffected by SR 49059 (*n =* 8, paired t-test, *t*(7) = 1.460, *P =* 0.1877).

### NMDAR-driven activity recruits rapid V1aR-dependent feedback to limit firing in vasopressin neurons

The results above indicate that pharmacological, exogenous application of VP efficiently modulated sAHP and spiking properties in VP neurons. We next aimed to determine whether similar effects could be evoked by endogenous, somatodendritically released VP^5^. We first tested whether an endogenous “tone” of extracellular VP, or activity-dependent somatodendritic release of VP during a burst of action potentials, could modulate sAHP and firing properties similarly to exogenously applied VP. To this end, we repeated the experiments above, but this time bath-applying a VP V1a receptor (V1aR) blocker (SR 49059, 1 µM). Interestingly, SR 49059 failed to affect the sAHP evoked by a fixed train (**Fig. 1E**) or by a single depolarizing step **(Supplementary Fig. 1B)**. Corresponding Ca^2+^ levels evoked by the trains were mostly unchanged (except for the decay tau, which was significantly higher (**Fig. 1F**)). Similarly, neither the number of evoked spikes (**Fig. 1G**) nor SFA adaptation **Fig. 1H**) were altered by the V1aR blocker. Together, these data argue against a substantial tonic V1aR-dependent modulation of sAHP or firing under these conditions and suggest that conventional train- or step-evoked activity is insufficient to recruit a detectable endogenous VP feedback signal. These results are consistent with our previous finding that spiking evoked by direct current depolarization failed to induce significant somatodendritic release of VP^32^.

We next tested whether a stronger, more physiologically relevant recruitment pattern could engage endogenous VP signaling. Based again on our prior work showing that NMDAR-driven, but not current step-driven, firing evokes somatodendritic VP release^32^, we next tested whether NMDAR-evoked spiking would induce VP-mediated modulation of the evoked sAHP. To maximize NMDAR availability and prevent activation of glutamate AMPA receptors, recordings were obtained in DNQX and low Mg^2+^ aCSF supplemented with glycine to reduce the Mg^2+^ pore block and facilitate Ca^2+^ influx through NMDAR. Glutamate uncaging at NMDAR (GluUC-NMDAR) targeted at VP somata consistently evoked a burst of action potentials. This NMDAR evoked firing displayed SFA approximately twice as slow compared to step depolarizations, and a slower repolarization to baseline that masked an underlying AHP. This was likely due to the slow diffusion kinetics of the uncaged glutamate which obscured the underlying AHP waveform. Notably, we found that bath-application of the V1aR blocker SR 49059 increased the number of NMDAR-evoked spikes while increasing the adaptation tau (**Fig. 2B,C**), indicating slower/weaker spike frequency adaptation. Importantly, these effects were not attributable to the modified uncaging solution itself. In low-Mg²⁺ aCSF containing glycine and caged glutamate, exogenous VP continued to enhance the sAHP, whereas SR 49059 remained without effect on sAHPs evoked by either conventional spike trains or depolarizing current steps (**Supplementary Fig. 2**). These findings indicate that the V1aR-dependent effect reflects the NMDAR-driven firing mode rather than the recording conditions per se. It appears NMDAR-driven activity recruits an endogenous V1aR-dependent inhibitory feedback signal that enhances adaptation and restrains spike output, consistent with rapid somatodendritic release of VP acting in an autocrine manner. They further indicate that this endogenous VP feedback is neither constitutively engaged nor recruited by all patterns of activity but instead is state- and activity-pattern dependent.

**Figure 2.**
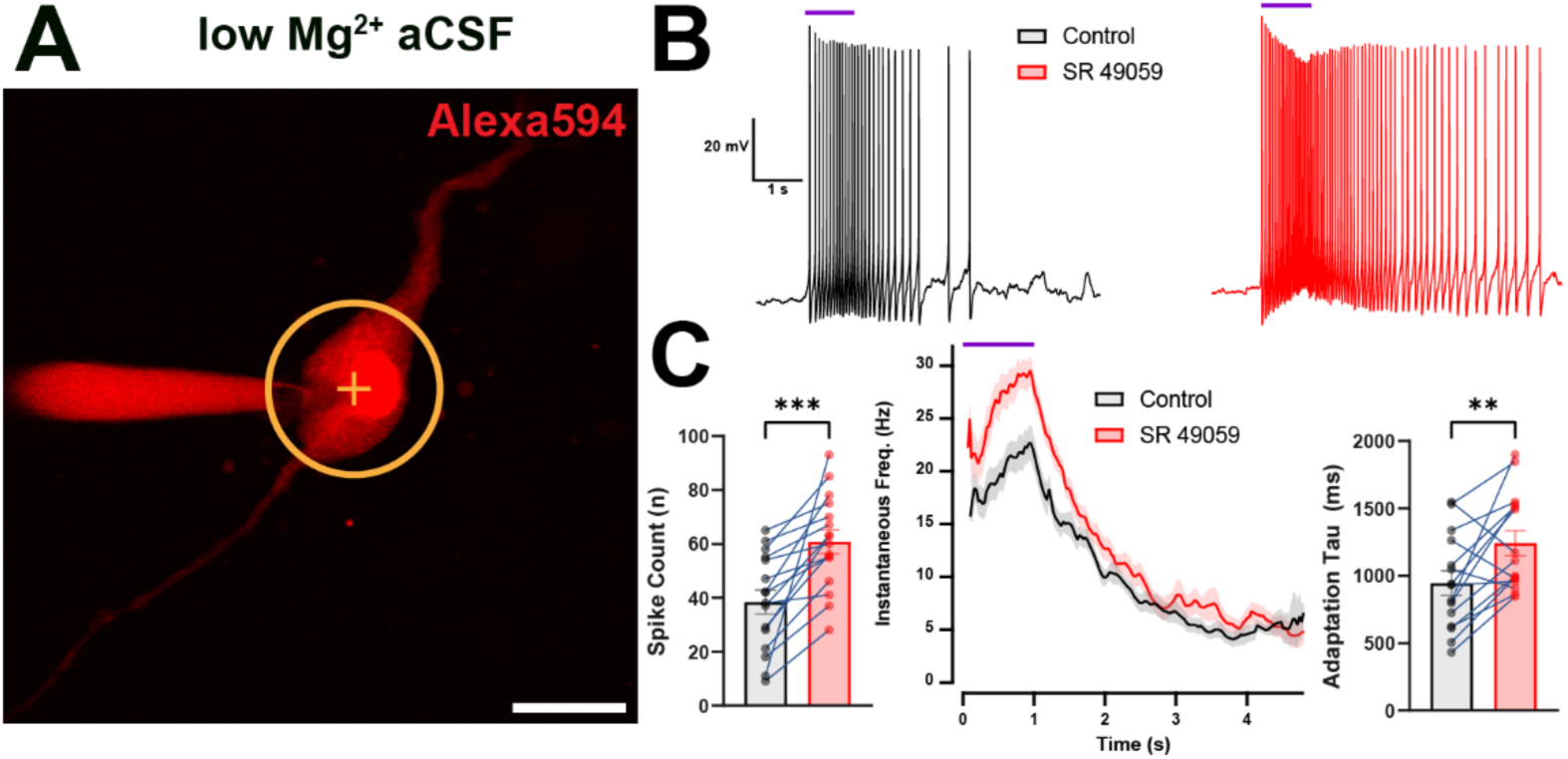
V1aR block enhances GluUC-NMDAR evoked spiking and reduced spike frequency adaptation. **(A)** Two-photon image of a patched VP neuron loaded with Alexa594 via the patch pipette in low Mg^2+^ aCSF. The target represents the area in which UV light was directed for GluUC-NMDAR. scale bar = 20 µm. **(B)** Representative example recording of a VP neuron responding to GluUC-NMDAR (1 s spiral, blue bar) before and during exogenous bath application of SR 49059. **(C)** gluUC-NMDAR summary data before and during SR 49059. (*left*) Number of spikes evoked in response to gluUC-NMDAR (*n =* 16, Wilcoxon test, *P =* 3.10E-05). (*middle*) Instantaneous frequency plotted as a function of time shows the trajectory of gluUC-NMDAR (blue bar) evoked spike frequency adaptation. (*right*) Adaptation taus are significantly higher during SR 49059 (*n =* 15, paired t-test, *t*(14) = 3.027, *P =* 0.0091).

### ER Ca²⁺ stores are required for sAHP-dependent adaptation during NMDAR-evoked spiking

Our previous work demonstrated that thapsigargin (TG) depletion of ER Ca^2+^ stores abolishes the sAHP in VP neurons^22^. Therefore, we used TG as a functional perturbation to test whether an ER store-dependent sAHP contributes to SFA during NMDAR-evoked firing. Slices were preincubated with TG (1 µM) for 1 h and remained exposed throughout the recordings. Consistent with our previous findings, TG-treated neurons displayed significantly inhibited sAHP amplitude and area (**Fig. 3A,B**). Ca^2+^ peak, area, and baseline fluorescence remain unchanged in this condition, although Ca^2+^ decay tau significantly increased (**Fig. 3C,D**). We interpret the prolonged decay as consistent with SERCA inhibition reducing ER Ca²⁺ reuptake capacity in these neurons. Additionally, TG increased spike count and adaptation tau during spiking evoked by depolarizing steps **(Supplementary Fig. 3A),** while sAHP amplitude, area, and index were all inhibited **(Supplementary Fig. 3B)**. Importantly, TG similarly increased spike output and adaptation tau during NMDAR-evoked firing (**Fig. 3E,F**). Together, these findings indicate that ER Ca²⁺ stores are required for full expression of the sAHP and contribute substantially to the adaptive restraint of firing during NMDAR-driven activity. In combination with the V1aR antagonist results, these data support a model in which NMDAR-evoked activity recruits a VP- and ER store-dependent inhibitory mechanism leading to the enhancement of the sAHP.

**Figure 3.**
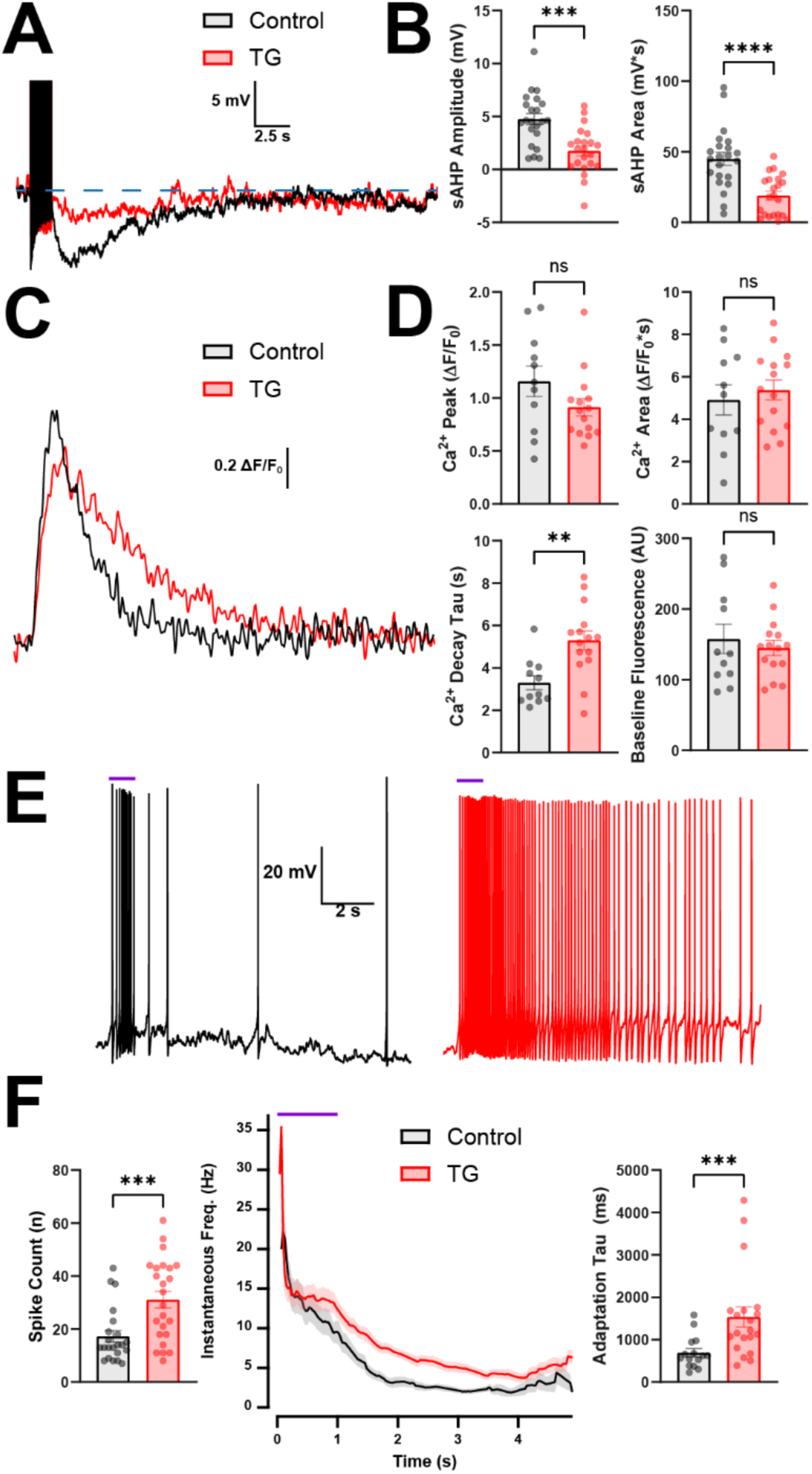
ER Ca^2+^ depletion enhances GluUC-NMDAR evoked spiking and reduced spike frequency adaptation. **(A)** Representative example of sAHPs from two different slice-matched cells under control conditions (black) and after 1+ hour of TG incubation (red). **(B)** TG incubation inhibits sAHP amplitude (Control *n =* 22 TG *n =* 22, paired t-test, *t*(42) = 4.284, *P =* 0.0001) and area (Control *n =* 22 TG *n =* 22, paired t-test, *t*(42) = 4.711, *P <* 0.0001). **(C)** Corresponding Ca^2+^ signals of cells in A. **(D)** TG incubation has no effect on Ca^2+^ peak (Control *n =* 11 TG *n =* 15, paired t-test, *t*(24) = 1.571, *P =* 0.1294), Ca^2+^ area (Control *n =* 11 TG *n =* 15, paired t-test, *t*(24) = 0.5686, *P =* 0.5749), or baseline fluorescence (Control *n =* 11 TG *n =* 15, paired t-test, *t*(24) = 3.371, *P =* 0.0025). Ca^2+^ decay tau significantly increased (Control *n =* 11 TG *n =* 15, paired t-test, *t*(24) = 0.5949, *P =* 0.5575). **(E)** Representative example of spiking evoked by gluUC-NMDAR (blue bar) under control conditions (black) and after 1+ hour of TG incubation (red). **(F)** Summary data of TG gluUC-NMDAR responses. TG incubation significantly increased spike count in response to gluUC-NMDAR (Control *n =* 22 TG *n =* 24, paired t-test, *t*(44) = 3.560, *P =* 0.0009) as well as adaptation tau (Control *n =* 15 TG *n =* 20, Mann-Whitney test, *P =* 0.0008).

### NMDAR activation transiently potentiates the sAHP through endogenous V1aR-dependent signaling

To more directly test whether endogenous VP-SDR modulates the sAHP, we paired a conventional sAHP assay with a preceding episode of NMDAR-driven spiking. sAHPs were evoked every 30 s by depolarizing pulse trains, and after two baseline trials, glutamate was uncaged at the recorded VP neuron 10 s before the third train to induce NMDAR-dependent firing and VP-SDR (**Fig. 4A**). This manipulation produced a robust but transient potentiation of the sAHP, reflected by increases in both amplitude and area that persisted across several subsequent sweeps (**Fig. 4B,C**). The potentiation was abolished by the V1aR antagonist SR49059, demonstrating that it depends on endogenous V1aR signaling (**Fig. 4B,C**). Notably, this effect was not accompanied by a corresponding V1aR-dependent enhancement of the train-evoked somatic Ca²⁺ transient (**Supplementary Fig. 4**), arguing that the sAHP potentiation is not simply secondary to a larger measured Ca²⁺ signal during the sAHP-evoking stimulus. These data therefore provide direct evidence that NMDAR-driven activity recruits an endogenous VP signal that transiently potentiates the sAHP, consistent with rapid autocrine somatodendritic release.

**Figure 4.**
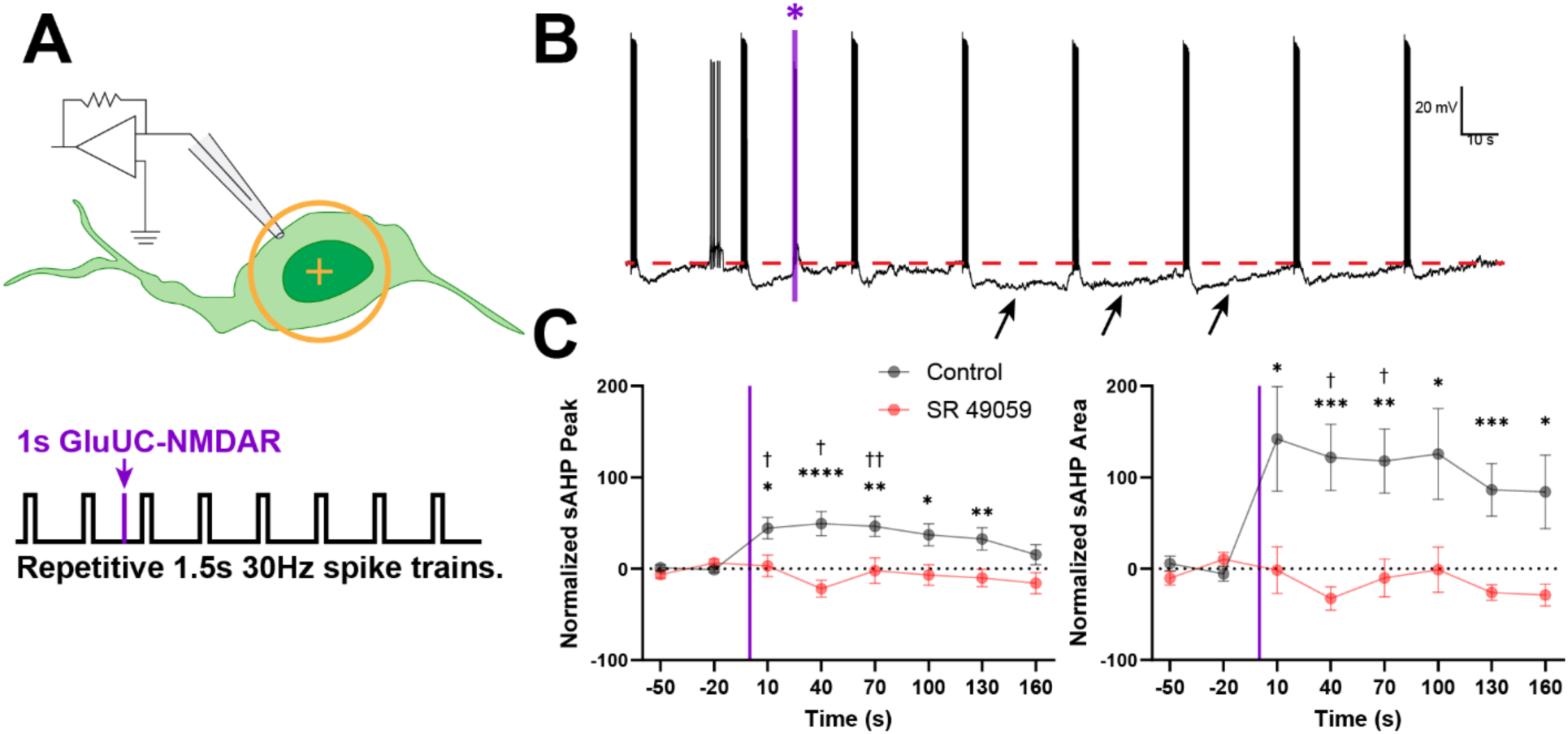
NMDAR-evoked somatodendritic release of VP (VP-SDR) transiently enhances sAHPs in an autocrine manner. **(A)** Graphic summarizing the experimental approach. VP neurons are patched and sAHPs are evoked every 30s *via* current injection. After two baseline measurements, gluUC-NMDAR on the recorded neuron (yellow target) occurs 10 s before the third sAHP stimulus. **(B)** Representative example traces of sAHPs during the protocol outlined in A. Vertical violet bar denotes gluUC-NMDAR stimulus. Note that the sAHP area enhances at sweeps 2-4 post gluUC-NMDAR (arrows). **(C)** Summary data for sAHP modulation in response to gluUC-NMDAR under control conditions (black) and in the presence of SR49059 (red). Normalized sAHP amplitude significantly increased. Peak: (*n =* 24, two way RM ANOVA; Time *F*(4.281, 184.1) *=* 2.802 *P =* 0.0243, Drug *F*(1, 43) *=* 12.59 *P =* 0.0010, Interaction *F*(4.281, 184.1) *=* 3.530 *P =* 0.0070). Area: (*n* = 24, two way RM ANOVA; Time *F*(3.566, 156.9) *=* 2.462 *P =* 0.0542, Drug *F*(1, 44) *=* 10.78 *P =* 0.0020, Interaction *F*(3.56, 156.9) *=* 2.812 *P =* 0.0326). Tukey multiple comparisons between drug conditions (*) and relative to baseline (†) are marked in the graphs.

### Paracrine VP signaling modulates the sAHP of neighboring VP neurons in a distance- and time-dependent manner

We previously showed that somatodendritically released VP from a single neuron can diffuse in the extracellular space and act in a paracrine manner to modulate the activity of nearby neurons^31^. We therefore asked whether VP-SDR from one VP neuron could modulate the sAHP of another, and whether this effect depended on intersomatic distance. To test this, we used the same sAHP assay described above in a patched VP neuron, but triggered gluUC-NMDAR in a neighboring VP neuron located at a known distance from the recorded cell (**Fig. 5A,B**). This design allowed us to examine the spatiotemporal profile of paracrine VP signaling while avoiding direct NMDAR activation of the neuron in which the sAHP was measured.

**Figure 5.**
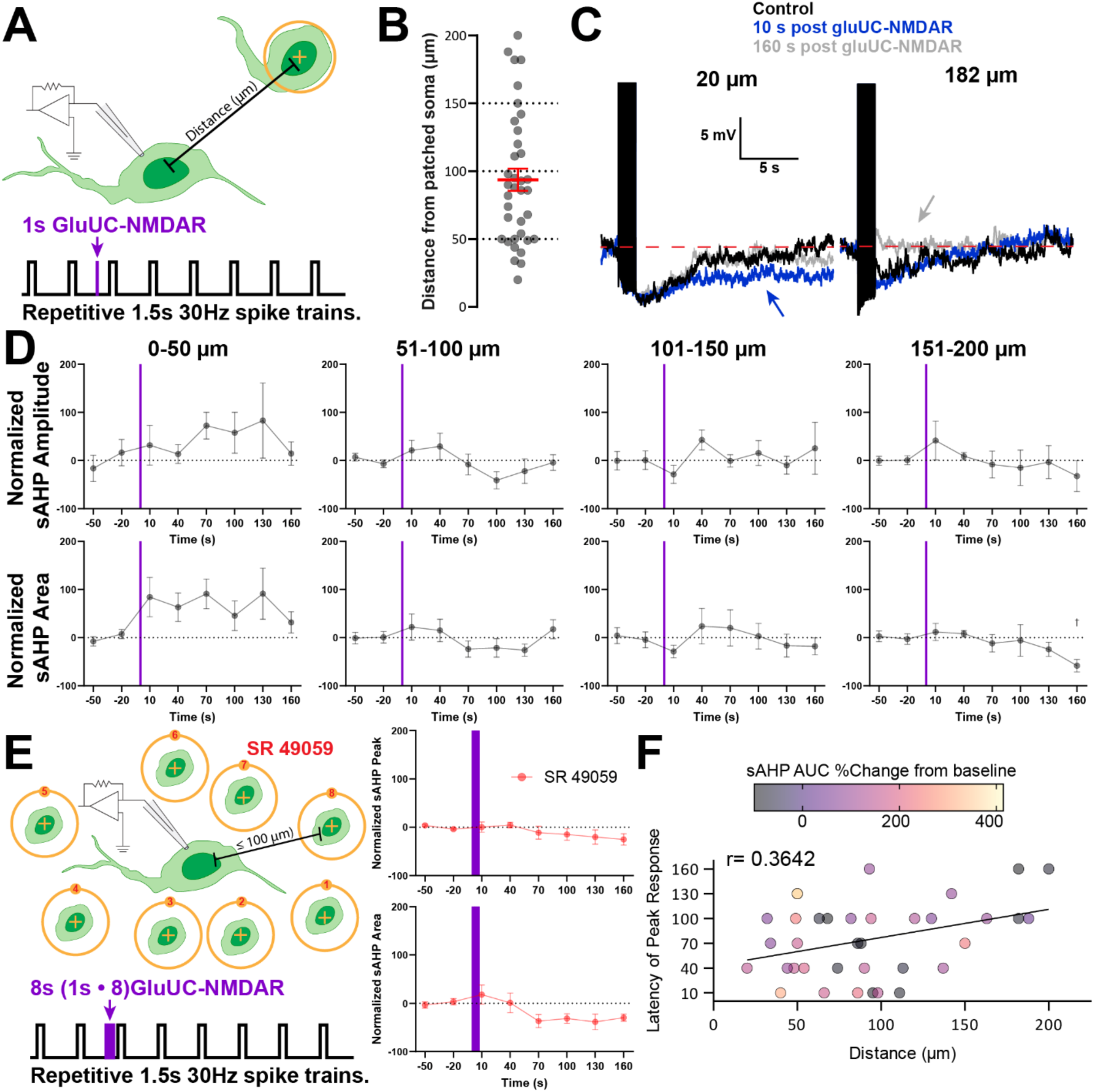
NMDAR-evoked somatodendritic release of VP (VP-SDR), acting in a paracrine manner, differentially modulates sAHP as a function of distance. **(A)** Graphic summarizing the experimental approach. VP neurons are patched and sAHPs are evoked every 30s *via* current injection. After two baseline measurements, gluUC-NMDAR on another neuron in the same focal plane (yellow target) occurs 10 s before the third sAHP stimulus. **(B)** Distance between patched and gluUC-NMDAR stimulated VP neuron for all trials. Mean distance was 93.7 ± 8.1 µm (n= 36 pairs). To evaluate VP-SDR’s paracrine effect, this data was binned by pair distance (dotted lines). **(C)** Example sAHPs from two different cells, one in which gluUC-NMDAR was in close proximity (*left*, 20 µm) and one that was far (*right*, 182 µm). For both trials, the sAHP at baseline, 10 s after gluUC-NMDAR, and 180 s after gluUC-NMDAR are superimposed. Note the close stimulation produces an enhancement after a short time and the long stimulation produces a delayed inhibition. **(D)** Normalized sAHP amplitude (*top*) and area (*bottom*) binned by distance. Graphs are separated for clarity, but two way RM ANOVA was performed with time and distance as the factors. Peak: (*n =* 36, two way RM ANOVA; Time *F*(3.545, 113.4) *=* 0.4323 *P =* 0.7624, Distance *F*(3, 32) *=* 1.070 *P =* 0.3757, Interaction *F*(10.64, 113.4) *=* 1.323 *P =* 0.2229). Area: (*n* = 36, two way RM ANOVA; Time *F*(4.260, 136.3) *=* 1.031 *P =* 0.3960, Distance *F*(3, 32) *=* 4.810 *P =* 0.0071, Interaction *F*(12.78, 136.3) *=* 1.273 *P =* 0.2375). **(E)** V1aR antagonism prevents sAHP modulation by gluUC-NMDAR. (*left*) Graphic summarizing the experimental approach. Eight neruons within 100 µm of the patched neuron were stimulated consecutively to robustly stimulate VP SDR. Other than the magnitude of the uncaging stimulus, sAHPs are evoked as before every 30s via current injection. (*right*) Summary data of pooled paracrine stimulation in the presence of SR 49059 (1 µM). Peak: (*n =* 7, one way RM ANOVA, *F*(2.476, 14.85) *=* 1.257, *P =* 0.3198). Area: (*n =* 7, one way RM ANOVA, *F*(2.5146, 15.27) *=* 2.365, *P =* 0.1181). **(F)** Latency of peak %Change in sAHP area relative to baseline plotted as a function of difference. There is a statistically significant positive correlation between distance and latency of peak response (*n =* 36, linear regression, *F*(1, 34) *=* 5.20, *r =* 0.3642, *P =* 0.0290). There is also a significant negative correlation between distance the peak %change values (*n =* 36, linear regression, *F*(1, 34) *=* 4.372, *r =* 0.3375, *P =* 0.0441) Points are colored with a heatmap and correspond to the magnitude of the %change, with a negative %change colored black.

Across 36 paired recordings spanning intersomatic distances from 20 to 200 µm, neighboring VP neurons exhibited qualitatively distinct responses depending on distance (**Fig. 5B,C**). At short distances, gluUC-NMDAR in a neighboring neuron enhanced the sAHP of the patched neuron, whereas at the greatest distances tested the response shifted toward a delayed inhibition (see representative examples in **Fig. 5C**).

Consistent with our previous data above, gluUC-NMDAR at neighboring VP neurons did not produce a clear distance-dependent modulation of the train-evoked somatic Ca²⁺ signal in the patched neuron (**Supplementary Fig. 5**). Thus, the bidirectional paracrine effects on sAHP are not readily explained by corresponding changes in the measured somatic Ca²⁺ transient. To analyze this systematically, recordings were grouped into 50-µm distance bins and sAHP magnitude was plotted over time before and after neighboring-neuron stimulation (**Fig. 5D**). This analysis revealed a significant distance-dependent effect on sAHP area, with robust enhancement at short distances, little or no net effect at intermediate distances, and a delayed inhibitory response emerging at the greatest distances tested (*n* = 36, two way RM ANOVA; Time *F*(4.260, 136.3) *=* 1.031 *P =* 0.3960, Distance *F*(3, 32) *=* 4.810 *P =* 0.0071, Interaction *F*(12.78, 136.3) *=* 1.273 *P =* 0.2375). Similar trends were observed for sAHP amplitude, although these did not reach significance (*n =* 36, two-way RM ANOVA; Time *F*(3.545, 113.4) *=* 0.4323 *P =* 0.7624, Distance *F*(3, 32) *=* 1.070 *P =* 0.3757, Interaction *F*(10.64, 113.4) *=* 1.323 *P =* 0.2229).

To test whether this paracrine modulation was mediated specifically by VP signaling, we repeated the experiment in the presence of the V1aR antagonist SR49059. In this control condition, gluUC-NMDAR was delivered sequentially to multiple neighboring VP neurons within 100 µm of the patched cell to maximize local VP release (**Fig. 5E**). Even under these conditions, SR49059 abolished the modulation of sAHP peak and area, indicating that the paracrine effect requires V1aR activation and arguing against mediation by other diffusible signals released during neuronal activation.

To further define the temporal organization of this paracrine response, we plotted for each recorded neuron the latency to peak sAHP modulation as a function of intersomatic distance, together with the magnitude and polarity of the response (**Fig. 5F**). This analysis revealed a significant relationship between distance and response latency (*r =* 0.3642, *P = 0.0290*), such that sAHP enhancements clustered at shorter distances and shorter latencies, whereas inhibitory responses predominated at greater distances and emerged more slowly. The progressively delayed onset of responses with increasing distance is consistent with passive extracellular diffusion contributing to the spread of paracrine VP signaling.

To further refine the spatiotemporal profile of paracrine VP signaling, we examined how the relationship between intersomatic distance versus sAHP modulation evolved over the full post-stimulation time course (**Supplementary Fig. 6**). We plotted regression slopes between these two variables at each timepoint post gluUC-NMDAR and implemented a cluster bootstrap that respected the repeated-measures structure of the data (bootstrapping neurons as clusters and retaining all time points per neuron). For each time point we calculated the regression slope of AUC versus distance and obtained 95% cluster-bootstrap confidence intervals. This analysis revealed an early distance-dependent enhancement at short range, a transient intermediate phase in which the spatial relationship weakened, and a later re-emergence of distance dependence dominated by inhibitory effects at greater distances. Together, these analyses indicate that paracrine VP signaling is dynamically organized over time, with early short-range enhancement giving way to a transient reduction in spatial structure before a later distance-dependent inhibitory phase emerges.

### Somatic, but not dendritic, VP signaling potentiates the sAHP in vasopressin neurons

Previous work has demonstrated that MNCs are highly compartmentalized both structurally and functionally^40–42^. We therefore asked whether VP modulation of the sAHP depends on the subcellular site of receptor activation. To address this, we used laser photolysis of a novel caged vasopressin (caged VP; see Methods) to selectively elevate local VP concentrations adjacent to either the soma or a primary dendrite of identified VP neurons. We validated specificity of caged VP by photoactivating it in the presence of “sniffer” cells plated onto the slice preparation (see Methods). Sniffer cells consisted of CHO cells transfected with a Ca^2+^ indicator and overexpressing either V1a receptors (sniffer_VP_) or oxytocin receptors (sniffer_OT_)^32,43,44^. Photoactivation of caged VP successfully activated 78.9% of sniffer_VP_ but failed to elicit any responses in sniffer_OT_ (**Supplementary Fig. 7**).

As in previous experiments, sAHPs were evoked every 30 s in a patched VP neuron, while VP was uncaged near either the soma or a primary dendrite of the recorded cell (**Fig. 6A**). Representative recordings from both compartments are shown in **Fig. 6B** and **6E**. Somatic VP uncaging produced a significant enhancement of both sAHP peak amplitude and area, whereas dendritic VP uncaging failed to significantly alter either measure (**Fig. 6C,D,F,G**). These findings indicate that VP modulation of the sAHP is compartment-specific and is effectively recruited at the soma, but not at dendritic sites.

**Figure 6.**
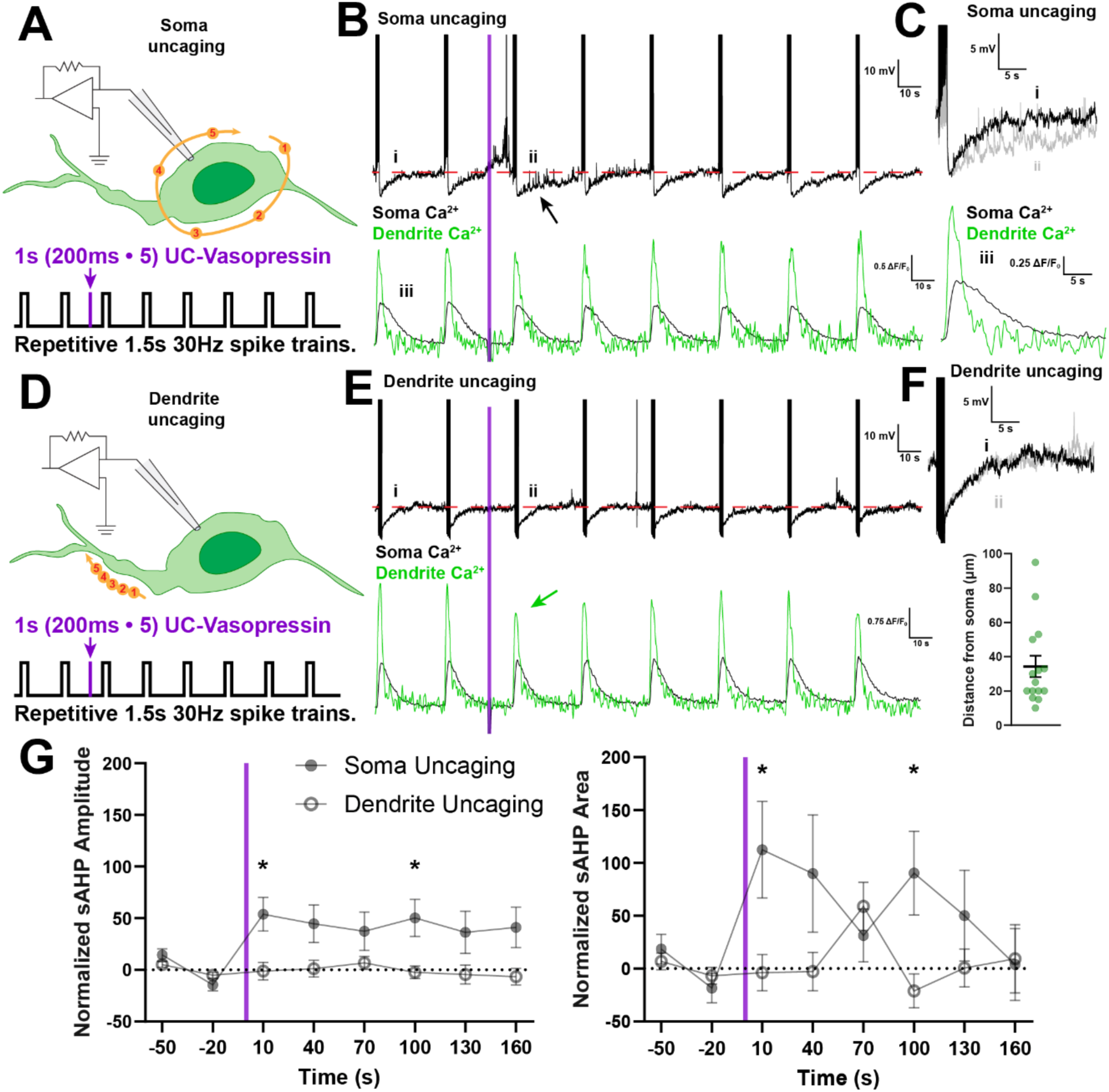
Somatic, but not dendritic, VP signaling potentiates the sAHP. **(A)** Graphic summarizing the experimental approach for uncaging at the soma. VP neurons are patched and sAHPs are evoked every 30s *via* current injection. After two baseline measurements, VP uncaging is triggered at points around the soma (yellow spots) 10 s before the third sAHP stimulus. **(B)** Representative example traces of sAHPs (*top*) and corresponding Ca^2+^ signal (*bottom*) when VP is uncaged at the soma. Vertical violet bar denotes caged VP stimulus. Note the sAHP enhancement after VP uncaging (arrow). **(C)** Zoomed examples of the traces in B to highlight important features. (*top*) AHP at baseline and immediately after caged VP superimposed on top of one another. Roman numerals correspond to data in B. Note the enhancement of the sAHP area. (*bottom*) Ca^2+^ signal from soma and dendrite of sweep 1. Note the differing kinetics of the Ca^2+^ signal between compartments. The somatic Ca^2+^ peak is shorter, but the overall signal decays more slowly compared to dendrite Ca^2+^. **(D)** Graphic summarizing the experimental approach. VP neurons are patched and sAHPs are evoked every 30s *via* current injection. After two baseline measurements, VP uncaging is triggered at points along the dendrite (yellow spots) 10 s before the third sAHP stimulus. **(E)** Representative example traces of sAHPs (*top*) and corresponding Ca^2+^ signal (*bottom*) when VP is uncaged at the dendrite. Vertical blue bar denotes caged VP stimulus. Note the decrease in dendritic Ca^2+^ immediately following a dendritic VP uncaging. **(F)** (*top*) Zoomed example of the electrophysiology trace in E. AHP at baseline and immediately after caged VP superimposed on top of one another. Roman numerals correspond to data in E. (*bottom*) Distance of all dendrite VP uncaging trials from the somatic membrane. **(G)** sAHP summary data of normalized sAHP amplitude and area when VP uncaging occurs at soma and dendrite. Two way RM ANOVA Tukey post hoc revealed uncaging compartment differences at 10 and 100 s post-stimulus for both normalized sAHP amplitude and area. Peak: (*n =* 36, two way RM ANOVA; Time *F*(3.366, 90.89) *=* 2.462 *P =* 0.0833, Uncaging Compartment *F*(1, 27) *=* 4.614 *P =* 0.0408, Interaction *F*(3.366, 90.89) *=* 2.462 *P =* 0.0608). Area: (*n* = 36, two way RM ANOVA; Time *F*(3.596, 86.31) *=* 1.559 *P =* 0.1975, Uncaging Compartment *F*(1, 24) *=* 2.851 *P =* 0.1043, Interaction *F*(3.596, 86.31) *=* 3.375 *P =* 0.0159). Significant post hoc differences are demarcated in the figure legend (\**P* < 0.05).

Although the representative recordings also illustrate compartmental differences in the associated Ca²⁺ signals, quantitative analysis showed that local VP uncaging did not produce corresponding changes in the train-evoked Ca²⁺ transient at either somatic or dendritic locations, despite elevating baseline Ca²⁺ in both compartments (**Supplementary Fig. 8**). Together, these findings indicate that VP-dependent potentiation of the sAHP is compartment-specific and preferentially linked to somatic signaling domains, consistent with a perisomatic locus for the relevant V1aR-sensitive machinery.

## DISCUSSION

Our findings identify the slow afterhyperpolarization (sAHP) as a key target through which somatodendritically released vasopressin (VP-SDR) regulates magnocellular VP neuron excitability. We show that VP enhances the integrated sAHP and strengthens spike frequency adaptation, and that endogenous VP released during NMDAR-driven activity recruits this inhibitory mechanism through rapid autocrine feedback. We further show that VP-SDR extends beyond the releasing neuron to modulate the sAHP of neighboring VP neurons in a distance- and time-dependent manner, and that this modulation is preferentially engaged at somatic rather than dendritic compartments. Together, these results support a model in which somatodendritic VP release couples intrinsic excitability to local spatiotemporal organization within the magnocellular VP network. We propose this mechanism to contribute to the generation of an appropriate VP population level response to cope with a homeostatic perturbation.

### Exogenous VP identifies the sAHP as a target of vasopressin signaling

Consistent with previous studies, we found that exogenously applied VP inhibited the firing activity of VP neurons. This was accompanied by an enhancement of the sAHP, mediated largely by a prolonged duration, delaying the membrane potential’s return to baseline. The VP-mediated enhancement of the sAHP occurred with a concomitant increase in the overall Ca^2+^ magnitude during the depolarizing step. This result is consistent with previous studies showing that VP not only invokes Ca^2+^ responses^37,38^, but elevates resting Ca^2+^ levels as well^45^. Moreover, we previously showed that that sAHP activation is almost entirely dependent on calcium availability in the cytosol^21^ and ER^22^. Thus, the concomitant increase in intracellular Ca²⁺ suggests a plausible mechanism by which VP could potentiate the sAHP, although a causal link remains to be directly established. Collectively, these results demonstrate that VP inhibits the firing activity of VP neurons, an effect mediated, at least in part, by potentiation of the sAHP.

### Activity-dependent autocrine VP signaling recruits transient sAHP feedback to restrain firing

To determine whether endogenously released VP produces an inhibitory effect similar to that observed with exogenous VP, we used an approach previously shown to efficiently evoke somatodendritic peptide release: NMDA receptor-evoked firing activity^32^. We found that focal glutamate uncaging under conditions favoring NMDAR activation reliably triggered a burst of action potentials in the targeted VP neuron. Importantly, V1aR blockade with SR49059 enhanced gluUC-NMDAR-evoked firing, increasing spike output and weakening spike frequency adaptation during the evoked burst (**Fig. 2**). These findings indicate that NMDAR-driven activity rapidly recruits endogenous VP signaling that feeds back onto the releasing neuron to restrain ongoing firing. Given that VP-SDR is strongest during NMDAR-evoked firing, compared to current injection, this specific mechanism may be engaged under certain circumstances such as during conditions of dehydration, when tight astrocyte wrappings have receded^46^ and extrasynaptic NMDARs become functionally more available^47,48^. This aligns with our observations that VP-sAHP modulation preferentially occurs at somatic compartments (**Fig. 6**).

Our previous work showed that depletion of ER Ca²⁺ stores with thapsigargin (TG) abolishes the sAHP in VP neurons^22^. Consistent with the contribution of this intrinsic inhibitory mechanism, TG produced a similar increase in NMDAR-evoked spike output and reduction in adaptation to that observed during V1aR blockade (**Fig. 3**). To test the involvement of the sAHP more directly, we repeatedly evoked sAHPs at fixed intervals before and after gluUC-NMDAR stimulation of the recorded neuron. NMDAR-driven activity produced a robust but transient increase in both sAHP amplitude and area, and this potentiation was completely prevented by SR49059 (**Fig. 4**). Thus, NMDAR-driven somatodendritic VP release rapidly engages autocrine V1aR signaling to enhance the sAHP. Together with the effects of TG and V1aR blockade on firing, these results strongly implicate sAHP potentiation as a key effector of VP-mediated autoinhibition, although they do not yet establish that this potentiation is necessary for the reduction in spike output.

The rapid onset of this feedback is consistent with the known ability of VP receptors to engage intracellular signaling on a timescale relevant to the evoked burst. In VP-sensitive supraoptic neurons, V1aR activation rapidly elevates intracellular Ca²⁺ through pathways involving extracellular Ca²⁺ entry, PLC-dependent signaling, and TG-sensitive intracellular ER stores^38,49,50^. Given the strong dependence of the sAHP on ER Ca²⁺ availability, a parsimonious model is that NMDAR-driven firing triggers somatodendritic VP release, which then activates pre-existing somatic V1aRs to enhance Ca²⁺-dependent coupling to the sAHP machinery and restrain continued firing. Magnocellular neurons exhibit several other rapid functional interactions between NMDARs and intrinsic or postsynaptic effectors, including A-type K⁺ currents^51^, GABA_A_ receptors^52^, and SK channels^53^, supporting the capacity of these neurons for fast local receptor–effector crosstalk. Whether V1aRs participate in a dedicated signaling complex with NMDARs or the sAHP machinery, however, remains unknown. Importantly, the absence of an SR49059 effect during conventional current-evoked trains is best interpreted as a failure of that activity pattern to recruit sufficient VP release, rather than as evidence against the sAHP as a downstream effector of VP-mediated autoinhibition.

Autocrine regulation of intrinsic afterpotentials has an important precedent in VP neurons. Dynorphin, which is co-packaged with VP in large dense-core vesicles, is released somatodendritically during phasic activity and acts through κ-opioid receptors to suppress the DAP and associated plateau potential that sustain burst firing^28,34,54,55^. Consistent with distinct temporal roles for these co-released peptides, V1a receptor blockade increases firing throughout the burst, whereas the effect of κ-opioid receptor blockade develops progressively as the burst proceeds^40^. Our findings extend this framework by identifying the sAHP as a complementary intrinsic target of VP signaling. Thus, dynorphin and VP may restrain phasic activity through distinct but convergent mechanisms: dynorphin reduces the depolarizing drive that sustains the burst, whereas VP strengthens the slow hyperpolarizing feedback that promotes spike frequency adaptation. Whether these pathways interact to shape burst duration remains to be tested directly.

### Paracrine VP signaling may shape asynchronous population output

At the population level, the functional demands placed on VP neurons differ fundamentally from those of oxytocin neurons. During lactation, oxytocin neurons are recruited into highly synchronized bursts that generate the large, pulsatile hormone release required to drive mammary smooth muscle contraction^56,57^. In contrast, effective VP secretion during an osmotic challenge depends on a more sustained and distributed population output, such that circulating VP reflects the integrated activity of many neurons firing asynchronously over time^14,58^. This organization is thought to help maintain continuous hormone availability while distributing secretory load across the population and avoiding fatigue of individual neurons. What remains unclear, however, is whether this asynchronous population behavior emerges largely from stochastic fluctuations in activity, or whether it is actively stabilized by local signaling interactions among VP neurons. Our findings raise the possibility that somatodendritic VP signaling contributes to this organization by locally shaping excitability across the population in space and time.

Our data demonstrate that NMDAR-driven somatodendritic VP release from a single neuron can alter the sAHP of a neighboring VP neuron, identifying the sAHP as an intrinsic target of endogenous paracrine VP signaling (Fig.5). This extends our previous work showing that somatodendritically released VP can diffuse through extracellular space and signal other neuronal populations^31,32^. Notably, this paracrine VP effect was organized in a spatiotemporal manner. We found that short-range signaling between VP neurons separated by 0-50 µm produced a relatively rapid potentiation of the sAHP, whereas little net modulation was observed at intermediate distances of 50–150 µm. At the greatest distances examined, 151–200 µm, the response shifted toward a delayed inhibition of the sAHP. The progressive increase in response latency with intersomatic distance is consistent with extracellular peptide diffusion contributing to the observed effects. A complementary time-resolved analysis further showed that the spatial pattern of paracrine VP action changes over time, progressing from early sAHP enhancement at short distances to delayed inhibition at longer distances.

The reversal in response polarity across distance is particularly intriguing. If extracellular diffusion simply produced a progressively lower concentration of VP acting through a single mechanism, one would expect the potentiation of the sAHP to diminish with distance, rather than reverse into inhibition. The transition from rapid sAHP enhancement at short range to delayed inhibition at greater distances therefore suggests that VP may engage distinct signaling mechanisms depending on the concentration and temporal profile experienced by the target neuron. Nearby neurons are likely exposed to a relatively rapid, high-concentration VP transient, whereas more distant ones may experience a lower-concentration but more prolonged signal.

Such differences in ligand concentration, rate of rise, and duration could promote distinct V1aR-dependent signaling pathways or receptor–effector coupling. Dose-dependent reversal of neuropeptide actions were previously reported in the magnocellular system, as low and high concentrations of oxytocin can exert opposing effects on neuronal excitability by engaging distinct intracellular signaling pathways (Gq vs.Gi, respectively)^59,60^. Alternatively, the delayed long-range response could involve indirect V1aR-dependent signaling through intervening neurons or glial cells^61^, or recruitment of another conductance that opposes the measured sAHP rather than direct inhibition of the same sAHP-generating machinery. Although V1aR blockade establishes that VP signaling is required for the paracrine response, it does not determine whether the short- and long-range effects arise through the same cellular pathway. Thus, the bidirectional response argues that paracrine VP signaling is not simply a graded consequence of diffusion but instead may engage qualitatively distinct mechanisms across spatial and temporal scales.

At the population level, these spatially structured paracrine effects raise the possibility that VP-SDR helps distribute excitability across the magnocellular network. Autocrine VP signaling would restrain the VP neurons undergoing strong activity, while rapid sAHP potentiation in nearby cells could reduce the probability of simultaneous recruitment within the same local cluster. In contrast, the delayed reduction in sAHP observed at greater distances could increase the relative excitability of more remote neurons. Such a pattern could generate spatial contrast across the VP population, limiting local synchrony while permitting activity to be redistributed across the population over time. This organization would be well suited to the physiological demands of the VP system, in which sustained hormone secretion depends on the integrated output of neurons firing in a distributed and largely asynchronous manner rather than through the highly synchronized bursts characteristic of oxytocin neurons^57,62^. By spreading secretory activity across the population, this mechanism could also help maintain continuous VP availability while reducing the burden placed on individual neurons during prolonged homeostatic challenges.

### Somatic VP signaling preferentially engages the sAHP

Our uncaging experiments further indicate that VP modulation of the sAHP is compartment-specific. Local VP elevation near the soma potentiated the sAHP, whereas comparable uncaging near a primary dendrite did not, despite evidence that V1aRs are present in both compartments^63^. These findings suggest that the relevant V1aR-sensitive signaling machinery is functionally concentrated within a somatic or perisomatic domain. This could reflect compartment-specific receptor–effector coupling, restricted localization of the sAHP-generating conductance, or differences in local Ca²⁺ signaling. Such compartmentalization is consistent with previous studies showing that somatic and dendritic compartments in magnocellular neurons are electrotonically decoupled^40,41^ and rely on distinct mechanisms to detect osmotic changes^42^. Together, our results suggest that geometric distance alone does not determine the efficacy of paracrine VP signaling with the route by which VP reaches the target cell and the extracellular microenvironment may be equally important.

### Concluding Remarks

Together, our findings identify the sAHP as a key intrinsic target of somatodendritic VP signaling in magnocellular neurons. Endogenous VP released during NMDAR-driven activity engages this mechanism through autocrine feedback to restrain firing and through paracrine signaling to shape the excitability of neighboring VP neurons in a distance- and time-dependent manner. The preferential recruitment of this response at somatic and perisomatic domains further indicates that the efficacy of VP signaling depends not only on peptide availability, but also on the spatial organization of receptor-effector coupling. These results therefore extend the role of the sAHP beyond simple spike-frequency adaptation, positioning it as a mechanism through which local neuropeptide release can link the activity of individual neurons to the organization of excitability across the magnocellular population. This framework may help explain how sustained and distributed VP output is maintained during prolonged homeostatic demand

## METHODS

### Animals

All experiments were approved by the Georgia State University Institutional Animal Care and Use Committee (IACUC) and performed in accordance with National Institutes of Health’s Guide for the care and use of laboratory animals. All experiments were performed in male and female young adult Wistar rats in the late peripubertal to adolescent developmental stage (150-300g) with transgenic expression of eGFP tagged to VP (eGFP-VP)^33^. These rats received *ad libitum* food and water and were housed on a 12:12 light-dark cycle.

### Ex vivo slice preparation

On the day of the experiment, rats were anesthetized with pentobarbital (50 mg kg^-1^, i.p.) and then transcardially perfused with 30-40 ml of ice-cold sucrose artificial cerebrospinal fluid (aCSF) solution. This sucrose aCSF solution contained (in mM): 200 sucrose, 2.5 KCl, 1 MgSO_4_, 26 NaHCO_3_, 1.25 NaH_2_PO_4_, 20 D-glucose, 0.4 ascorbic acid, and 2.0 CaCl_2_, pH 7.2, 300-305 mOsmol l^-1^. The animal was then rapidly decapitated, and the brain was subsequently removed, mounted in the chamber of a vibrotome (Leica VT1200S, Leica Microsystems), with superglue and submerged into sucrose aCSF bubbled constantly with 95% O_2_/5% CO_2_. Slices were cut at 250 μm thickness, bisected at the midline to separate the bilateral SONs into their own separate slice, and placed in a holding chamber containing standard aCSF (in mM): 119 NaCl, 2.5 KCl, 1 MgSO_4_, 26 NaHCO_3_, 1.25 NaH_2_PO_4_, 20 D-glucose, 0.4 ascorbic acid, 2 CaCl_2_, and Na^+^ Pyruvate pH 7.2, 300-305 mOsmol l^-1^ bubbled with 95% O_2_/5% CO_2_. Slices rested at 32 °C *via* water bath for 20 minutes before transfer to room temperature for a minimum of 40 minutes before any recording.

### Whole cell patch clamp

Slices were placed in the chamber of an upright 2 photon microscope (Bruker Nano Inc.) and perfused constantly by aCSF warmed to 32 °C at a flow rate of 2 ml min^-1^. Cells were acquired in whole cell current clamp configuration using a MultiClamp 700B and digitized with a Digidata 1550b (Molecular Devices). Electrodes were pulled using a Flaming Brown horizontal puller (Sutter Instruments) from borosilicate capillaries (1.5 mm OD, 1.17 mm ID, 75 mm length; Warner Instruments) with a 3-7 MΩ resistance and filled with internal solution (in mM): 135 KMeSO_4_, 8 KCl, 10 HEPES, 2 Mg-ATP, 0.3 Na-GTP, 6 phosphocreatine, as well as Alexa 594 to visualize the patched neuron, and 0.05 fluo-5F for Ca^2+^ imaging (7.2 pH; 285-292 mosmol (kg H_2_O)^-1^. Estimated liquid junction potential was 10 mV and not corrected. Once we achieved whole cell configuration, we allowed the cell to rest for 5 minutes to ensure stability of the patch and equilibration between internal solution and cytosol. We injected negative holding current (0-85 pA) to bring the cell to a baseline resting membrane potential of -55 mV. We evoked sAHPs using two different protocols to evaluate different aspects of the sAHP, including a pulse train with fixed spike count and a single 2.5s depolarizing step (see Experimental design and statistical analyses section for details). In experiments where the sAHP was evaluated over time (Fig. 4-6, Supplementary Figure 3), two baseline sAHPs were recorded to ensure stable sAHP generation. This was followed by an uncaging stimulus 10 s preceding the third sAHP. A total of 8 sAHPs were measured in each trial with 30 s between each one. Data was acquired at 10 kHz. Cells that displayed a shift in series resistance that exceeded 20 MΩ or a 25% change from the start of the recording were discarded.

### Two-photon Ca^2+^ imaging

All fluorescence imaging was performed using a tunable Mai Tai laser (MKS Spectra-Physics) set to 860 nm on a Bruker two-photon microscope running Prairie View software. Emitted fluorescence was separated using a 565-nm dichroic mirror, directing signal to 525 ± 30 nm (green) and 595 ± 33 nm (red) emission filters positioned in front of two photomultiplier tubes (PMTs). All imaging data was acquired using a 40x Olympus objective (LUMPLFLN40XW). eGFP-VP neurons and fluo-5F signals were collected in the green channel, while Alexa 594 was detected in the red channel. Ca^2+^ imaging acquisition was performed simultaneously with whole cell patch clamp *via* fluo-5F dialyzed intracellularly *via* patch pipette (0.05 mM). Ca^2+^ waveforms were acquired online using Prairie View’s brightness over time (BOT) functionality. Inert Alexa 594 was used as a reference channel to assist in targeting uncaging stimuli as well as monitor photobleaching. Ca^2+^ imaging data was generally acquired at 12 Hz, except in cases where we were capturing somatic and dendritic Ca^2+^ simultaneously. This necessitated a slower sampling rate that ranged from 5-12 Hz. Additionally in caged VP experiments some experiments did not have both a dendrite and soma in focus. In this scenario, only the compartment that was uncaged was imaged.

### Photolytic uncaging

Uncaging experiments were performed simultaneously with patch clamp and Ca^2+^ imaging within Prairie View utilizing Marked Points functionality to target light to specific regions of interest within the slice. We photolytically activated caged compounds using with a 405 nm Helios laser (30µm diameter spiral, 1 s; 3 mW). Experiments utilizing MNI-caged-glutamate (caged glu) required modified aCSF composition. Previous work in our group demonstrated that NMDAR activation triggers robust SDR relative to a simple depolarizing current injection^32^. Therefore, we isolated and potentiated NMDAR activation by excluding AP5, reducing MgSO_4_ to 0.1 mM, and adding 0.1 mM glycine to the aCSF (low-Mg^2+^ aCSF). We refer to this approach as glutamate uncaging at NMDARs (gluUC-NMDAR).

Caged vasopressin (caged VP, 1 µM) experiments necessitated an alternative uncaging protocol for two reasons. First, the caged VP compound properties required a higher 405 nm intensity to break the caged compared to caged glu. Second, we compared responses when uncaging VP at somas versus dendrites. Given the dendrites’ smaller surface area, we wanted to standardize VP uncaging between these two compartments as much as possible. Thus, caged VP was activated by point uncaging at 5 points equally spaced in a circle around the soma or at 5 equally spaced points in a 10 µm line parallel to the dendrite (5 points for 200 ms each, 1 s total; 7 mW). Uncaging occurred directly adjacent to the plasma membrane to minimize phototoxicity. See diagrams in Figure 6A or Supplementary Figure 4A.

### Sniffer Cells

Chinese hamster ovary (CHO) sniffer cells designed for detection of endogenous neuropeptide release^64^ were generated and utilized for to validate caged VP specificity **(Supplementary Fig. 7)**. These procedures have been described previously^32,43^. Briefly, CHO cells overexpressing human oxytocin receptor (OTR)s or vasopressin 1a receptor (V1aR), sniffer_OT_ and sniffer_VP_ respectively, were grown and seeded on 30 mm petri dishes. 12-18 hours before experimentation, sniffer_OT_ and sniffer_VP_ dishes were transfected with a solution of 124 µl sterile MilliQ water, 8.2 µl Fugene HD reagent (Promega, Madison, WI, USA) and 2.4 µl of a red shifted fluorescent Ca^2+^ indicator plasmid (R-GECO; GenScript, Piscataway, NJ, USA) into 30 mm dishes containing 2.5 mL of culture media at ∼80% confluency. On the day of the experiment, sniffer cell monolayers were detached from the dish with trypsin, resuspended in aCSF, and transferred directly onto a brain slice. Brain slices embedded with sniffer cells were perfused with aCSF containing caged VP and stimulated with a 300 ms 405 nm point of light. After each trial, a 10 µM bolus of VP or OT (corresponding to sniffer_VP_ or sniffer_OT_ respectively) was injected into the bath as a positive control to confirm sniffer cell viability. Cells that failed to respond to this bolus were discarded.

### Experimental design and statistical analyses

We operationally define the sAHP as the apamin-insensitive component of the afterpotential following a train of spikes until it returns to resting membrane potential. Therefore, baseline sAHPs were pharmacologically isolated under control conditions with 0.1 µM apamin to block the SK component of the AHP (mAHP)^12,35^, 5 mM CsCl to block the slow depolarizing afterpotential (sDAP), which overlaps in time course with the AHP^34,65^, as well as 10 µM DNQX, 40 µM AP5, and 100 µM Px to block AMPA, NMDA, and GABA receptors respectively. sAHPs were evoked using two separate protocols. The first is a pulse train consisting of 30 spikes at 20 Hz with a 5 ms pulse width was used to measure sAHP properties themselves; the fixed spike count better controls for total Ca^2+^ entering the cell, an important consideration for this Ca^2+^-dependent afterpotential^21,66,67^. The other protocol was a 2.5 second depolarizing 50 pA current step to evaluate how the sAHP impacts firing rate and spike frequency adaptation (SFA).

Electrophysiological data was analyzed using custom code (M.K. Kirchner) in MATLAB (The Mathworks Inc.) incorporating abfload.m to read .abf pClamp files (originally written by Harald Hentschke). sAHP amplitude is the peak amplitude of the sAHP relative to the baseline membrane potential (mV). sAHP area is the definite integral of the 10 s interval immediately after spike termination bounded by the signal and the baseline membrane potential (mV*ms). For experiments where we measured sAHPs in response to a single depolarizing step. sAHP amplitude and area are expressed as absolute values for clarity. Instantaneous frequency was calculated by taking the reciprocal of the time interval between spike events and generating a cumulative average across time. Adaptation taus were calculated by fitting a single exponential to the plot of a cell’s instantaneous frequency vs. time and calculating the time constant of this fit^39^. In experiments where we measured multiple sAHPs over time, we normalized amplitude and area in response to an average of two baseline sAHP measurements.

Ca^2+^ imaging data was analyzed from fluorescence waveform data acquired online with brightness over time (BOT) functionality in PrairieView. Region of interest was Fluorescence data was normalized to a 1 second baseline fluorescence measurement and expressed as the relative change in fluorescence to baseline (ΔF/F_0_). A 1 s gaussian blur filter was applied to the entire Ca^2+^ signal. Ca^2+^ peak is maximal change in Ca^2+^ relative to baseline (ΔF/F_0_). Ca^2+^ area is the definite integral of the signal interval from spike initiation timepoint (*t*_0_) to *t*_0_+20 s bounded by the signal and y=0 (ΔF/F_0_*s). Ca^2+^ decay tau is the time constant of a single exponential fit to the decay of Ca^2+^ signal from Ca^2+^ peak to *t*_0_+20 s. Baseline fluorescence is averaged intensity of the Ca^2+^ signal at baseline (AU). In experiments where we measured multiple sAHPs over time, we evaluated whether baseline Ca^2+^ changed in response to a stimulus. In these cases, we evaluated baseline fluorescence normalized to the average of the first two baseline fluorescence measurements (AU).

Statistical analysis was performed in Prism (GraphPad Inc.). Data was tested for normality using D’Agostino-Pearson omnibus and homogeneity of variance. If the data met these conditions, an appropriate parametric test was performed. Otherwise, an equivalent non-parametric test was performed instead. Each experiment included a minimum of four animals. n values represent the number of cells. No more than one cell per slice was evaluated in experiments where pharmacological manipulation was the independent variable. In TG experiments where the same neurons were unable to be evaluated before and after TG-mediated ER Ca^2+^ depletion, neurons were slice matched in almost all cases to minimize variability. In experiments where the sAHP was evaluated over time (Fig. 4-6, Supplementary Figure 3), the two baseline measurements were averaged together and a 2-Way RM ANOVA was performed on this data, however plots show both baseline values. For Ca^2+^ decay taus, often not every single trial could be reliably fit to a single exponential fit and these points were excluded from analysis. We therefore performed a mixed-effects analysis on RM Ca^2+^ decay tau data.

Bootstrap samples were generated by resampling the %Change in sAHP area paracrine dataset, not individual measurements. In each iteration, we selected N neurons with replacement and retained the entire set of %Change in sAHP area values for all time points from each selected neuron. 5000 iterations were performed. This approach ensured that the repeated-measures structure was preserved, time points were never mixed, and sampling variability was estimated at the neuron level. Plots were generated in Prism. Example traces were generated in Igor Pro 9 (Wavemetrics Inc.). Graphics were generated in Adobe Illustrator (Adobe Inc.).

### Reagents

10 μM 6,7-dinitroquinoxaline-2,3-dione (DNQX; HelloBio)

40 μM 2R)-amino-5-phosphonovaleric acid (AP5; HelloBio)

100 μM picrotoxin (Px; HelloBio)

0.1 µM apamin (apamin; Tocris Bioscience)

25 µM MNI-caged-glutamate (caged glu; Tocris Bioscience)

0.1 µM [Arg^8^]-Vasopressin (VP; Millipore Sigma)

1 µM SR 49059 (SR 49059; Tocris Bioscience)

1 μM thapsigargin (TG; Tocris Bioscience)

1 µM caged vasopressin (caged VP; gift from Ismail Ahmed)

50 µM fluo-5F, pentapotassium salt (fluo-5F; Invitrogen)

25 µM Alexa Fluor 594 Hydrazide (Alexa 594; Invitrogen)

### Resources

eGFP-VP rat (Yoichi Ueta)

MatLab abf reader (Harald Hentschke (2025) abfload.m)

(https://www.mathworks.com/matlabcentral/fileexchange/6190-abfload), MATLAB Central File Exchange.)

## Funding

M.K.K. NIH R00HL168434

J.E.S. NIH R01HL162575

I.A.H. Burroughs Wellcome Fund - CASI Award

---

**Supplementary Figure 1.**
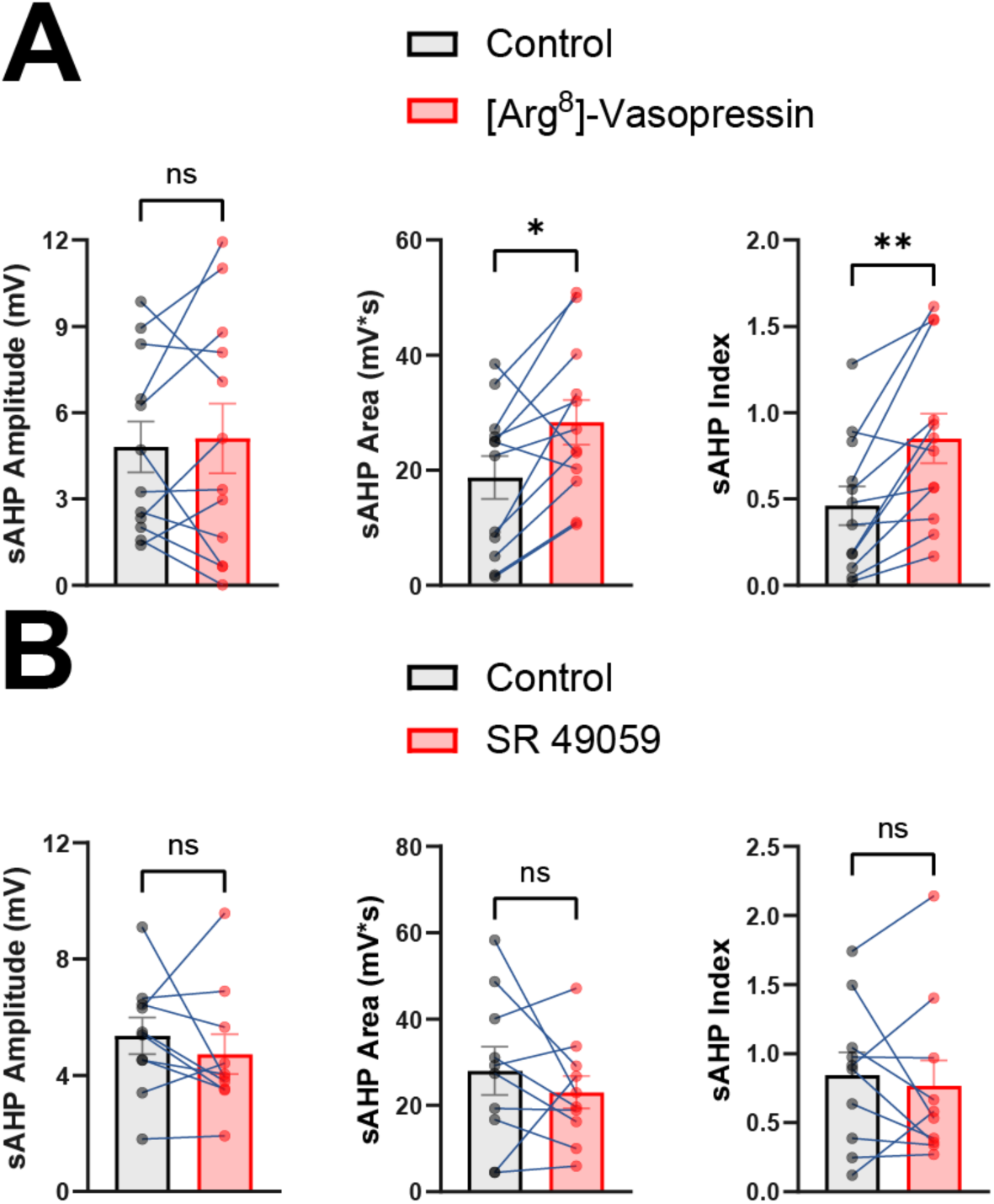
VP enhances sAHPs evoked by depolarizing current steps. **(A)** [Arg^8^]-Vasopressin enhances area (*n =* 12, paired t-test, *t*(11) = 2.945, *P =* 0.0133) and sAHP Index (*n =* 12, paired t-test, *t*(11) = 4.019, *P =* 0.0020), but not amplitude (*n =* 12, paired t-test, *t*(11) = 0.3869, *P =* 0.7062) of sAHPs evoked by a single 50 pA depolarizing current step. **(B)** SR 49059 does not affect amplitude (*n =* 10, paired t-test, *t*(9) = 0.9288, *P =* 0.3773), area (*n =* 10, paired t-test, *t*(9) = 0.9885, *P =* 0.3487), or sAHP index (*n =* 10, paired t-test, *t*(9) = 0.5404, *P =* 0.6020) of sAHPs evoked by a single 50 pA depolarizing current step.

**Supplementary Figure 2.**
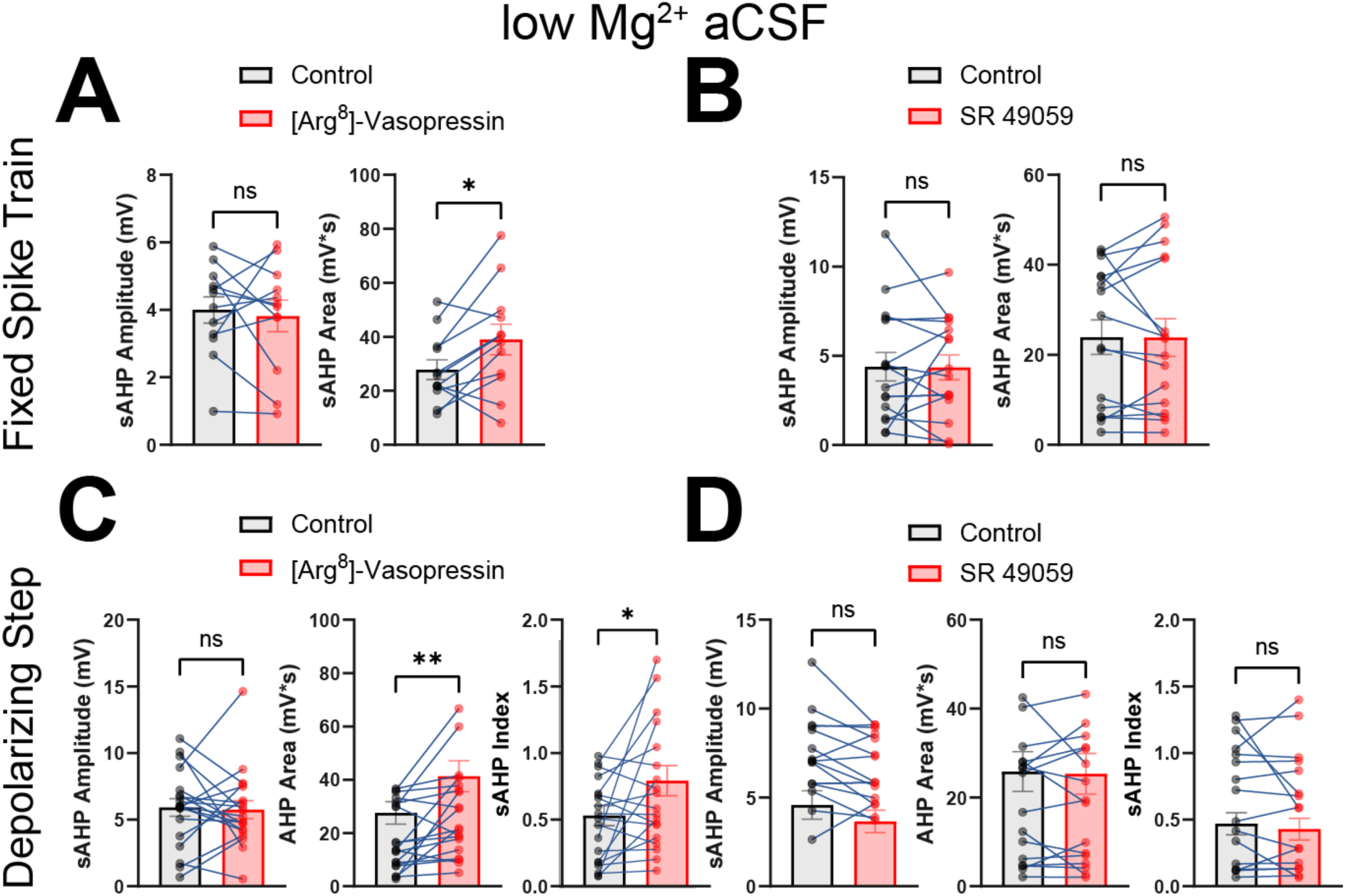
sAHPs evoked by depolarizing current steps in the presence of low Mg^2+^ aCSF. **(A)** sAHPs evoked by fixed spike trains in low Mg^2+^ aCSF and caged glu before and during exogenous bath application of [Arg^8^]-Vasopressin. sAHPs demonstrate significantly increased sAHP area (*n =* 12, paired t-test, *t*(11) = 2.731, *P =* 0.0195), but not amplitude (*n =* 12, paired t-test, *t*(11) = 0.4242, *P =* 0.6796) during [Arg^8^]-Vasopressin. **(B)** sAHPs evoked by pulse trains in low Mg^2+^ aCSF and caged glu before and during exogenous bath application of SR 49059. sAHPs demonstrate no significant differences in SR 49059. sAHP amplitude (*n =* 16, paired t-test, *t*(15) = 0.0746, *P =* 0.9415), sAHP area sAHP area (*n =* 16, paired t-test, *t*(15) = 0.0406, *P =* 0.9682). **(C)** sAHPs evoked by a 50 pA depolarizing step in low Mg^2+^ aCSF and caged glu before and during exogenous bath application of [Arg^8^]-Vasopressin. sAHPs demonstrate significantly increased sAHP area (*n =* 19, paired t-test, *t*(18) = 3.063, *P =* 0.0067) and sAHP index (*n =* 19, paired t-test, *t*(18) = 2.400, *P =* 0.0274), but not amplitude (*n =* 19, paired t-test, *t*(18) = 0.2132, *P =* 0.8336) in the presence of [Arg^8^]-Vasopressin. **(D)** sAHPs evoked by pulse trains in low Mg^2+^ aCSF and caged glu before and during exogenous bath application of SR 49059. sAHPs demonstrate no significant differences in SR 49059. sAHP amplitude (*n =* 16, paired t-test, *t*(15) = 1.968, *P =* 0.0678), sAHP area (*n =* 16, paired t-test, *t*(15) = 2.515, *P =* 0.8147), sAHP index (*n =* 16, Wilcoxon test, *P =* 0.9799).

**Supplementary Figure 3.**
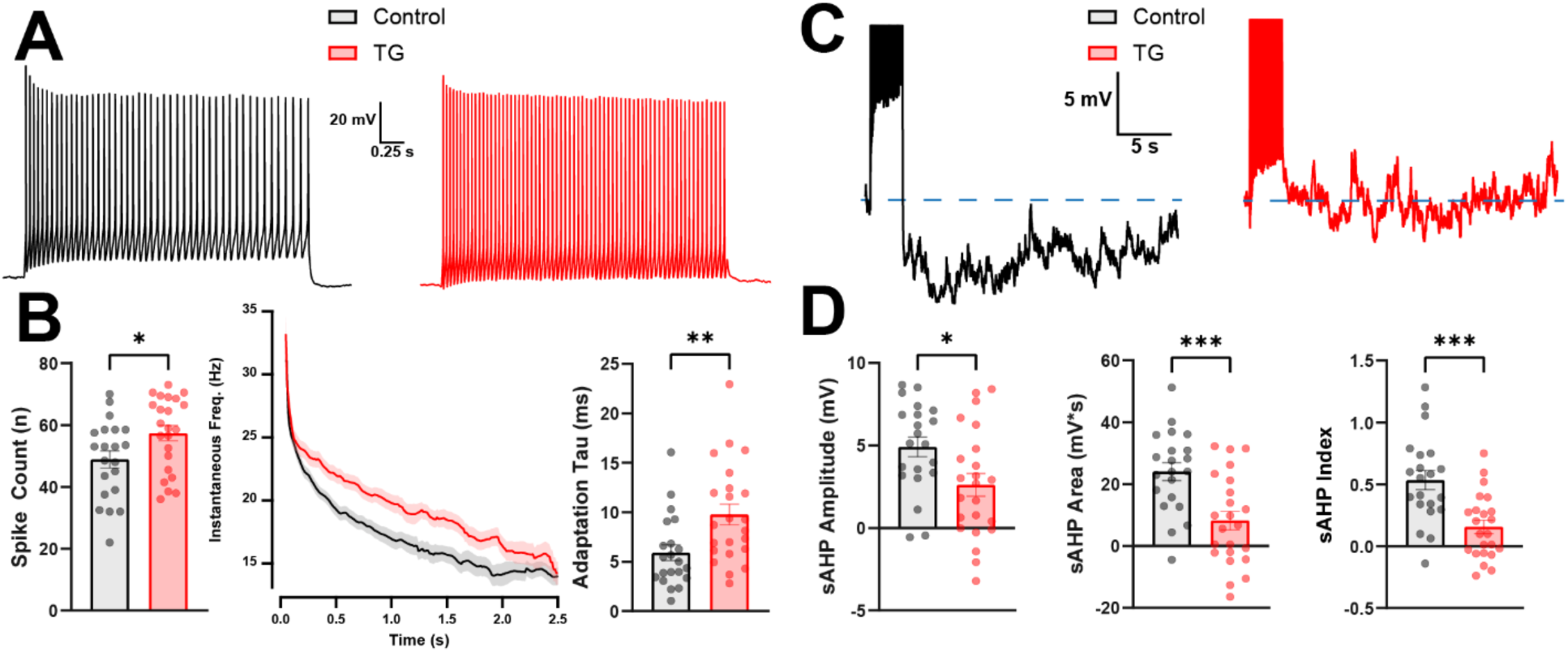
TG inhibits SFA and sAHPs when evoked by depolarizing current step. **(A)** Representative example traces of spiking from two slice-matched VP neurons in response to a single current step under control conditions or after 1h+ TG incubation. **(B)** Summary data of single step TG data. Spike count was significantly higher in TG compared to controls (Control *n =* 21 TG *n =* 23, paired t-test, *t*(42) = 2.285, *P =* 0.0275). Adaptation taus were longer in TG compared to controls (Control *n =* 21 TG *n =* 23, Mann-Whitney test, *P =* 0.0032). **(C)** sAHPs of the same traces from panel A scaled to visualize the sAHPs. **(D)** sAHPs are significantly inhibited in VP neurons incubated in 1h+ TG. sAHP amplitude (Control *n =* 21 TG *n =* 23, paired t-test, *t*(42) = 2.515, *P =* 0.0158); sAHP area (Control *n =* 21 TG *n =* 23, paired t-test, *t*(42) = 3.783, *P =* 0.0005); sAHP index (Control *n =* 21 TG *n =* 23, paired t-test, *t*(42) = 4.075, *P =* 0.0002).

**Supplementary Figure 4.**
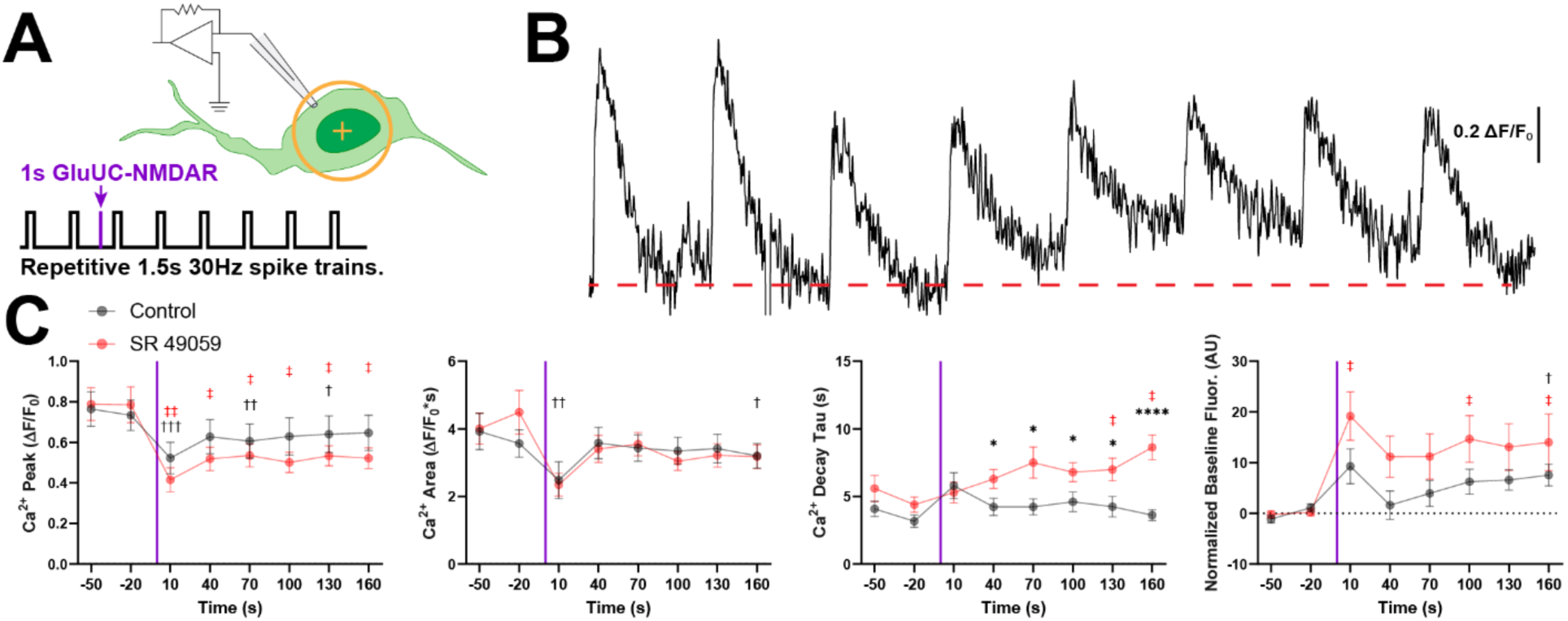
V1aR-dependent potentiation of the sAHP occurs without a corresponding enhancement in train-evoked somatic Ca²⁺ transients. **(A)** Graphic summarizing the experimental approach. VP neurons are patched and sAHPs are evoked every 30s *via* current injection. After two baseline measurements, gluUC-NMDAR on the recorded neuron (yellow target) occurs 10 s before the third sAHP stimulus. **(B)** Ca^2+^ response corresponding to the same recording in figure 4B. **(C)** Summary Ca^2+^ data of V1aR dependent potentiation of sAHPs. Ca^2+^ Peak: (*n =* 25, two way RM ANOVA; Time *F*(2.866, 65.93) *=* 15.75 *P <* 0.0001, Drug *F*(1, 23) *=* 0.9048 *P =* 0.3514, Interaction *F*(2.866, 65.93) *=* 1.684 *P =* 0.1809). Ca^2+^ Area: (*n* = 25, two way RM ANOVA; Time *F*(3.315, 76.25) *=* 8.116 *P <* 0.0001, Drug *F*(1, 23) *=* 0.0041 *P =* 0.9493, Interaction *F*(3.315, 76.25) *=* 0.6605 *P =* 0.5936). Ca^2+^ Decay Tau: (*n =* 25, mixed effects analysis; Time *F*(3.975, 82.15) *=* 1.671 *P =* 0.1648, Drug *F*(1, 23) *=* 6.377 *P =* 0.0189, Interaction *F*(3.975, 82.15) *=* 3.052 *P =* 0.0216). Ca^2+^ Normalized Baseline Fluorescence: (*n* = 25, two way RM ANOVA; Time *F*(1.894, 43.56) *=* 4.815 *P =* 0.0142, Drug *F*(1, 23) *=* 1.298 *P =* 0.2664, Interaction *F*(1.894, 43.56) *=* 1.742 *P =* 0.1886). Tukey multiple comparisons between drug conditions (*) and relative to baseline (†) are marked in the graphs.

**Supplementary Figure 5.**
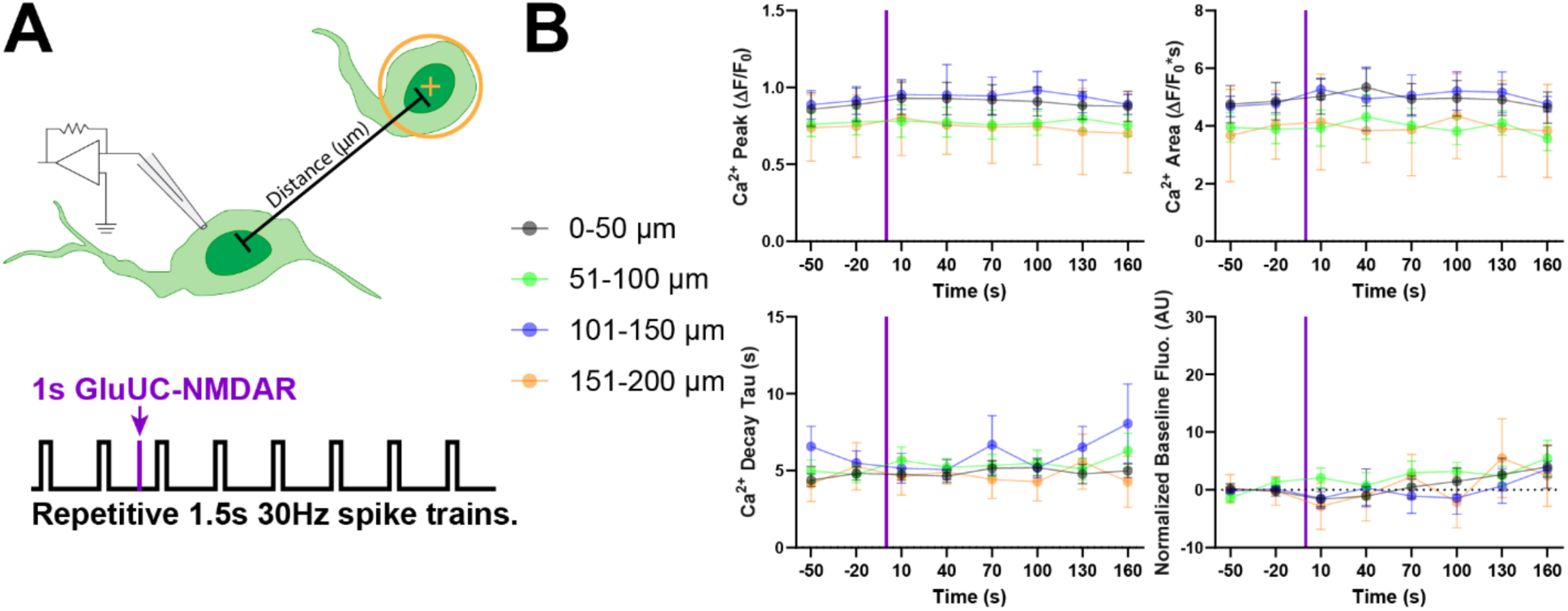
Paracrine VP signaling modulates the sAHP without corresponding distance-dependent changes in somatic Ca²⁺ transients. **(A)** Graphic summarizing the experimental approach. VP neurons are patched and sAHPs are evoked every 30s *via* current injection in the presence of SR49059. After two baseline measurements, gluUC-NMDAR on the recorded neuron (yellow target) occurs 10s before the third sAHP stimulus. **(B)** Corresponding Ca^2+^ signal data to paracrine VP SDR presented in Figure 5. Two-Way RM ANOVA revealed no significant differences except for in time of normalized baseline fluorescence. Ca^2+^ Peak: (*n =* 34, two way RM ANOVA; Time *F*(3.119, 93.56) *=* 1.658 *P =* 0.1795, Distance *F*(3, 30) *=* 0.5987 *P =* 0.6208, Interaction *F*(9.356, 93.56) *=* 0.4314 *P =* 0.9199). Ca^2+^ Area: (*n* = 34, two way RM ANOVA; Time *F*(3.599, 108) *=* 1.285 *P =* 0.2818, Distance *F*(3, 30) *=* 0.7635 *P =* 0.5235, Interaction *F*(10.80, 108) *=* 0.6605 *P =* 0.5936). Ca^2+^ Decay Tau: (*n =* 34, mixed effects analysis; Time *F*(2.718, 81.53) *=* 1.207 *P =* 0.3113, Distance *F*(3, 30) *=* 0.5929 *P =* 0.6245, Interaction *F*(8.153, 81.53) *=* 0.9231 *P =* 0.5033). Ca^2+^ Normalized Baseline: (*n* = 34, two way RM ANOVA; Time *F*(2.875, 86.25) *=* 3.663 *P =* 0.0167, Distance *F*(3, 30) *=* 0.2734 *P =* 0.8441, Interaction *F*(8.625, 86.25) *=* 0.6129 *P =* 0.7764).

**Supplementary Figure 6.**
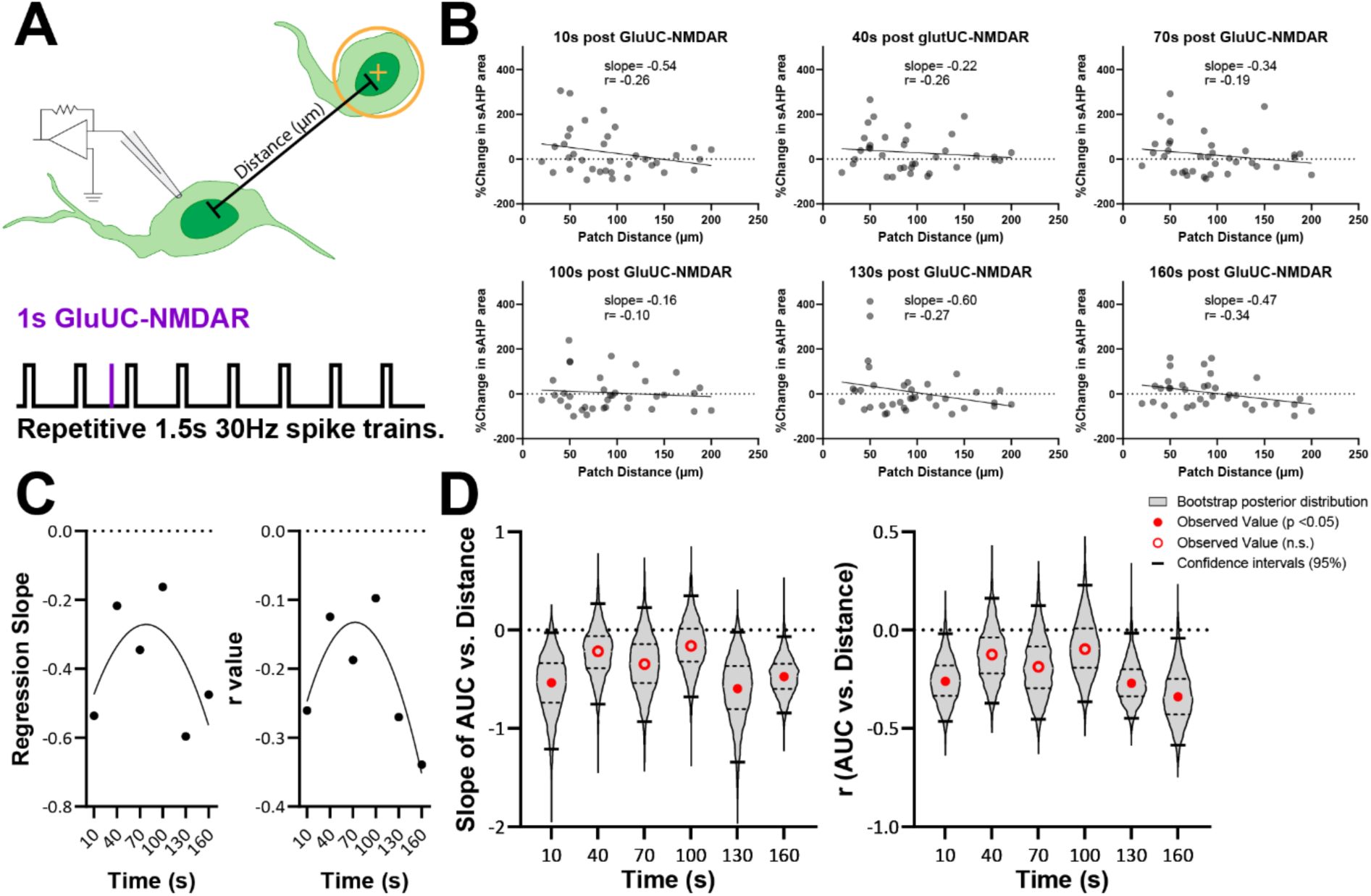
The relationship between distance and paracrine sAHP modulation changes over time. **(A)** Graphic summarizing the experimental approach. VP neurons are patched and sAHPs are evoked every 30s *via* current injection. After two baseline measurements, gluUC-NMDAR on another neuron in the same focal plane (yellow target) occurs 10 s before the third sAHP stimulus. **(B)** Plots of %change in sAHP area as a function of patch distance at each time point following a paracrine VP-SDR evocation. Each dataset is fitted with a linear regression. **(C)** regression slopes and r values calculated in panel B plotted as a function of time and fitted with a quadratic equation. Regression slope (*n =* 6, quadratic fit, *df =* 3, *R^2^ =* 0.415) and r value (*n =* 6, quadratic fit, *df =* 3, *R^2^ =* 0.772). **(D)** Bootstrap analysis of %Change in sAHP area vs. distance for slope of linear regression and r values. Note the transient shallowing of the slope and r value at the 40 s, 70 s, and 100 s timepoints, which recovers at 130 s and 160 s. Violin plots represent the bootstrap distribution, red dots represent the values of the observed values for regression slope and r value from panel C, and horizontal bars represent the confidence intervals. Note the upper confidence intervals reverse polarity at 40 s, 70 s, and 100 s for both slope of AUC and r. Slopes (10 s *slope =* -0.537, 95% bootstrap CI [-1.21, -0.03] *P >* 0.05; 40 s *slope =* -0.2171, 95% bootstrap CI [-0.75, 0.27] *P <* 0.05; 70 s *slope =* -0.345, 95% bootstrap CI [-0.93, 0.23] *P <* 0.05; 100 s *slope =* -0.162, 95% bootstrap CI [-0.68, 0.35] *P <* 0.05; 130 s *slope =* -0.596, 95% bootstrap CI [-1.34, -0.02] *P >* 0.05; 160 s *slope =* -0.475, 95% bootstrap CI [-0.84, -0.07] *P >* 0.05). r values (10 s *r =* -0.260, 95% bootstrap CI [-0.46, -0.02] *P >* 0.05; 40 s *r =* -0.125, 95% bootstrap CI [-0.37, 0.16] *P <* 0.05; 70 s *r =* -0.187, 95% bootstrap CI [-0.45, 0.12] *P <* 0.05; 100 s *r =* -0.097, 95% bootstrap CI [-0.36, 0.23] *P <* 0.05; 130 s *r =* -0.270, 95% bootstrap CI [-0.45, -0.02] *P >* 0.05; 160 s *r =* -0.339, 95% bootstrap CI [-0.59, -0.04] *P >* 0.05).

**Supplementary Figure 7.**
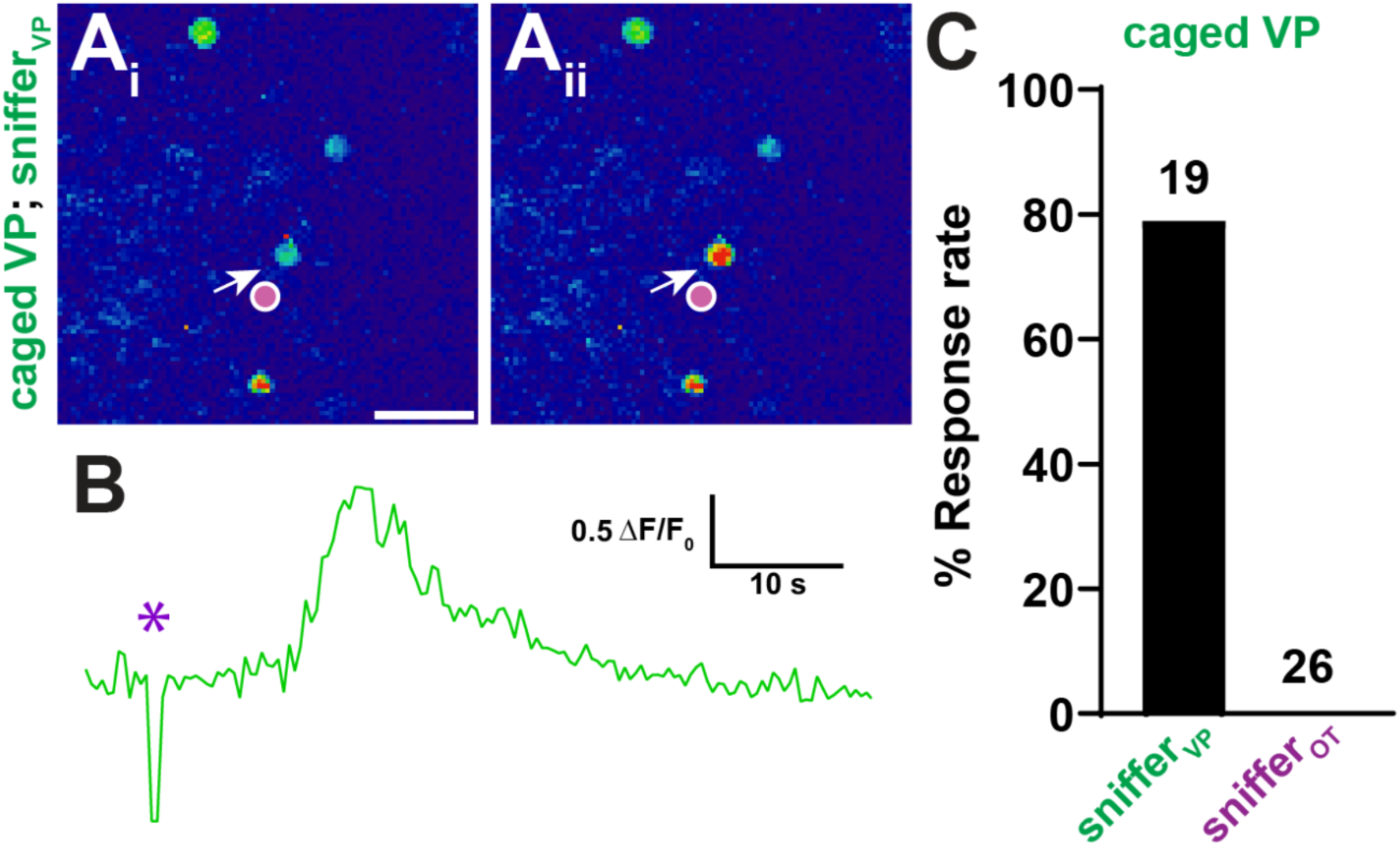
Photolysis of caged VP selectively activates V1aR-expressing sniffer cells. **(A)** Representative fluorescence images of V1aR-expressing sniffer cells (sniffer_VP_) transfected with the Ca²⁺ indicator R-GECO and imaged in the presence of caged VP. Before photolysis (i), caged VP produced no detectable activation of the indicated cell (arrow). A 300-ms pulse of 405-nm light targeted to the indicated region (purple circle) evoked a robust increase in intracellular Ca²⁺ fluorescence (ii). Scale bar, 50 µm. (B) Corresponding R-GECO Ca²⁺ trace from the cell shown in (A). The purple asterisk indicates the timing of the 405-nm photolysis pulse. (C) Response rates to photolysis of caged VP in V1aR-expressing sniffer cells (sniffer_VP_) and oxytocin receptor-expressing sniffer cells (sniffer_OT_). Photolysis of caged VP evoked Ca²⁺ responses in approximately 80% of snifferVP cells, whereas no responses were detected in sniffer_OT_ cells. Numbers above the bars indicate the number of cells tested in each group

**Supplementary Figure 8.**
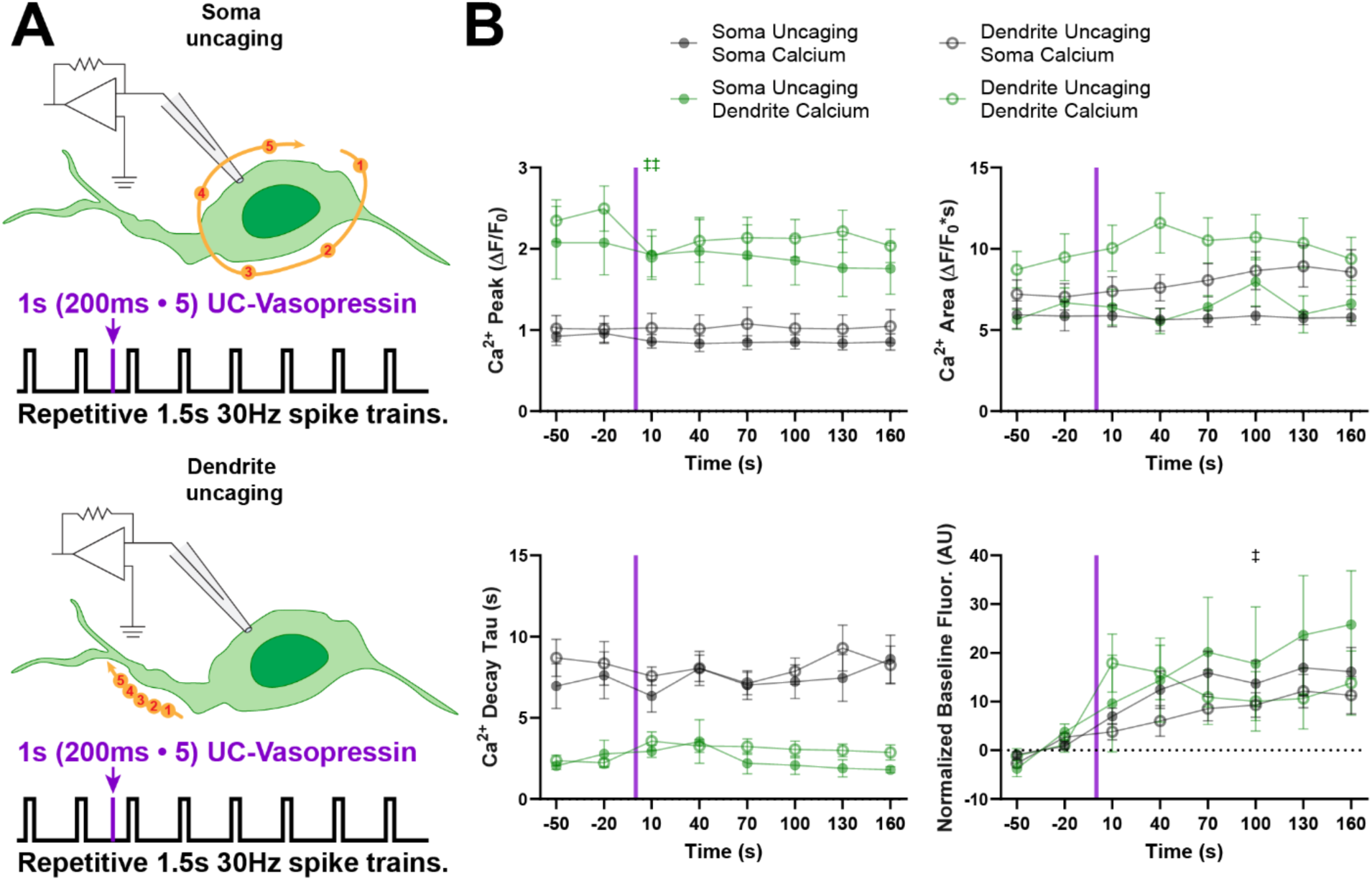
Somatic and dendritic Ca²⁺ signals exhibit distinct kinetics but are minimally altered by local VP uncaging. **(A)** Graphic summarizing the experimental approach for uncaging at the soma. VP neurons are patched and sAHPs are evoked every 30s *via* current injection. After two baseline measurements, VP uncaging is triggered at points around the soma (yellow spots) 10 s before the third sAHP stimulus. **(B)** Summary Ca^2+^ data during sAHP evocation. Two way RM ANOVA with Tukey post hoc testing revealed no significant differences were observed between uncaging soma and dendrite within the same compartment, however there are significant differences in the Ca^2+^ waveform between soma and dendrites in both Ca^2+^ peak and Ca^2+^ decay tau. Multiple comparisons are omitted for clarity, please see *Supplementary Table 1.* Peak: (*n =* 41, two way RM ANOVA; Time *F*(2.500, 92.49) *=* 3.988 *P =* 0.0148, Compartment *F*(3, 37) *=* 8.890 *P =* 0.0001, Interaction *F*(7.499, 92.49) *=* 2.025 *P =* 0.0557). Area: (*n* = 42, two way RM ANOVA; Time *F*(3.273, 124.4) *=* 1.723 *P =* 0.1611, Compartment *F*(3, 38) *=* 3.567 *P =* 0.0228, Interaction *F*(9.819, 124.4) *=* 1.404 *P =* 0.1872). Adaptation Tau: (*n =* 40, mixed effects analysis; Time *F*(3.744, 127.9) *=* 0.5881 *P =* 0.6608, Compartment *F*(3, 36) *=* 16.83 *P <* 0.0001, Interaction *F*(11.23, 127.9) *=* 0.8603 *P =* 0.5827). Baseline: (*n* = 41, two way RM ANOVA; Time *F*(2.969, 109.9) *=* 9.889 *P <* 0.0001, Compartment *F*(3, 37) *=* 0.4908 *P =* 0.6908, Interaction *F*(8.908, 109.9) *=* 1.255 *P =* 0.2701)

**Supplementary Table 1.** Statistical summary of Ca^2+^ multiple comparisons from Figure 6H.

| Time Point | Baseline | 10 s | 40 s | 70 s | 100 s | 130 s | 160 s |
| --- | --- | --- | --- | --- | --- | --- | --- |
| **Ca<sup>2+</sup> Peak** |  |  |  |  |  |  |  |
| Soma UC; Soma Ca<sup>2+</sup> vs. Soma UC; Dendrite Ca<sup>2+</sup> | 0.1372 | 0.0537 | 0.1272 | 0.1162 | 0.0682 | 0.1518 | 0.1209 |
| Soma UC; Soma Ca<sup>2+</sup> vs. Dendrite UC; Soma Ca<sup>2+</sup> | 0.9697 | 0.7903 | 0.7586 | 0.7179 | 0.8295 | 0.7922 | 0.8473 |
| Soma UC; Soma Ca<sup>2+</sup> vs. Dendrite UC; Dendrite Ca<sup>2+</sup> | 0.0004 | 0.0049 | 0.0016 | 0.0009 | 0.0008 | 0.0014 | 0.001 |
| Soma UC; Dendrite Ca<sup>2+</sup> vs. Dendrite UC; Soma Ca<sup>2+</sup> | 0.1801 | 0.1221 | 0.2322 | 0.2736 | 0.1533 | 0.2986 | 0.2957 |
| Soma UC; Dendrite Ca<sup>2+</sup> vs. Dendrite UC; Dendrite Ca<sup>2+</sup> | 0.897 | >0.9999 | 0.9936 | 0.9625 | 0.9497 | 0.8221 | 0.9736 |
| Dendrite UC; Soma Ca<sup>2+</sup> vs. Dendrite UC; Dendrite Ca<sup>2+</sup> | 0.001 | 0.0438 | 0.0122 | 0.0175 | 0.0097 | 0.0091 | 0.0324 |
| **Ca<sup>2+</sup> Area** |  |  |  |  |  |  |  |
| Soma UC; Soma Ca<sup>2+</sup> vs. Soma UC; Dendrite Ca<sup>2+</sup> | 0.9947 | 0.9766 | 0.9997 | 0.8456 | 0.5914 | 0.997 | 0.8905 |
| Soma UC; Soma Ca<sup>2+</sup> vs. Dendrite UC; Soma Ca<sup>2+</sup> | 0.7286 | 0.5365 | 0.2927 | 0.21 | 0.1842 | 0.1313 | 0.2787 |
| Soma UC; Soma Ca<sup>2+</sup> vs. Dendrite UC; Dendrite Ca<sup>2+</sup> | 0.1906 | 0.0681 | 0.0371 | 0.0232 | 0.0237 | 0.0486 | 0.0884 |
| Soma UC; Dendrite Ca<sup>2+</sup> vs. Dendrite UC; Soma Ca<sup>2+</sup> | 0.8149 | 0.8955 | 0.3018 | 0.5591 | 0.983 | 0.3424 | 0.6915 |
| Soma UC; Dendrite Ca<sup>2+</sup> vs. Dendrite UC; Dendrite Ca<sup>2+</sup> | 0.2251 | 0.2165 | 0.0365 | 0.0746 | 0.5581 | 0.1384 | 0.3965 |
| Dendrite UC; Soma Ca<sup>2+</sup> vs. Dendrite UC; Dendrite Ca<sup>2+</sup> | 0.561 | 0.4047 | 0.2354 | 0.505 | 0.6773 | 0.892 | 0.973 |
| **Ca<sup>2+</sup> Decay Tau** |  |  |  |  |  |  |  |
| Soma UC; Soma Ca<sup>2+</sup> vs. Soma UC; Dendrite Ca<sup>2+</sup> | 0.0114 | 0.0237 | 0.1034 | 0.0033 | 0.0031 | 0.0154 | 0.0044 |
| Soma UC; Soma Ca<sup>2+</sup> vs. Dendrite UC; Soma Ca<sup>2+</sup> | 0.8828 | 0.7127 | >0.9999 | 0.9997 | 0.9614 | 0.8012 | 0.9973 |
| Soma UC; Soma Ca<sup>2+</sup> vs. Dendrite UC; Dendrite Ca<sup>2+</sup> | 0.0221 | 0.1165 | 0.0049 | 0.009 | 0.0129 | 0.0503 | 0.0135 |
| Soma UC; Dendrite Ca<sup>2+</sup> vs. Dendrite UC; Soma Ca<sup>2+</sup> | 0.002 | <0.0001 | 0.0881 | 0.0016 | 0.0003 | 0.0028 | 0.0022 |
| Soma UC; Dendrite Ca<sup>2+</sup> vs. Dendrite UC; Dendrite Ca<sup>2+</sup> | 0.8768 | 0.5778 | 0.9969 | 0.5984 | 0.5888 | 0.3884 | 0.1886 |
| Dendrite UC; Soma Ca<sup>2+</sup> vs. Dendrite UC; Dendrite Ca<sup>2+</sup> | 0.0035 | 0.0004 | 0.001 | 0.0021 | 0.001 | 0.0083 | 0.0063 |
| **Norm. Ca<sup>2+</sup> Baseline** |  |  |  |  |  |  |  |
| Soma UC; Soma Ca<sup>2+</sup> vs. Soma UC; Dendrite Ca<sup>2+</sup> | - | 0.964 | 0.9979 | 0.9858 | 0.9892 | 0.9606 | 0.8686 |
| Soma UC; Soma Ca<sup>2+</sup> vs. Dendrite UC; Soma Ca<sup>2+</sup> | - | 0.9787 | 0.6527 | 0.5826 | 0.8801 | 0.9223 | 0.8914 |
| Soma UC; Soma Ca<sup>2+</sup> vs. Dendrite UC; Dendrite Ca<sup>2+</sup> | - | 0.2463 | 0.809 | 0.991 | 0.9993 | 0.9648 | 0.9975 |
| Soma UC; Dendrite Ca<sup>2+</sup> vs. Dendrite UC; Soma Ca<sup>2+</sup> | - | 0.9361 | 0.8008 | 0.7529 | 0.889 | 0.8027 | 0.6272 |
| Soma UC; Dendrite Ca<sup>2+</sup> vs. Dendrite UC; Dendrite Ca<sup>2+</sup> | - | 0.8658 | 0.9554 | 0.9545 | 0.9802 | 0.8487 | 0.8148 |
| Dendrite UC; Soma Ca<sup>2+</sup> vs. Dendrite UC; Dendrite Ca<sup>2+</sup> | - | 0.1868 | 0.2656 | 0.8433 | 0.9509 | >0.9999 | 0.9669 |

